# A Unifying Thermodynamic Model for Phase Separation and Aging of Biopolymers

**DOI:** 10.64898/2026.02.21.707146

**Authors:** Jasper J. Michels, Joana Caria, Edward A. Lemke

## Abstract

Protein condensates that form via phase separation typically become more viscous over time and can harden in a process referred to as “molecular aging”. Several mechanisms have been identified for this phenomenon. Of these, the ones involving enhanced β-sheet or -strand interactions are of pathological relevance since they have been associated with neurodegeneration. Although there is much understanding of biopolymer phase behavior, an inclusive thermodynamic framework that unifies phase separation and β-sheet-based aging is lacking. We present a time-dependent, multi-component extension of associating polymer theory that describes phase separation and aging of an intrinsically disordered protein (IDP) capable of associating through local, reversible folding. The model shows how the Second Law of Thermodynamics applies throughout, whether phase separation precedes and encourages aging or, vice versa, whether the increase in “stickiness” during aging drives phase separation. Our calculations show how the time-dependence of the average valency of associating sites determines the aging kinetics and the development of viscoelastic properties of a biocondensate. The agreement between our calculations and the change in dynamics of condensates of ‘perfect repeat’ analogues of nucleoporin-98 not only validates the theory but also identifies these Nup98 variants as model systems for studying aging.

## 1. Introduction

Biomolecular condensates are ubiquitous in eukaryotic cells, fulfilling a plethora of biological functions.^[1], [2]^ Many biomolecular condensates contain intrinsically disordered proteins (IDPs) and typically form via phase separation from the cyto- or nucleoplasm. Due to the fact that IDPs (and proteins in general) are associative heteropolymers, their condensates are microscopically heterogeneous^[3]^ and may transition between different material states, that identify as viscoelastic liquids or solids.^[4]^ Most biocondensates have the tendency to ‘age’ or ‘mature’, during which a natively high fluidity slowly transitions into a more solid-like state, usually through local conformational changes and increased multi-valent interactions. An important class of neuronal IDPs ages via enhanced β-strand-based interactions between their (prion-like) low-complexity domains (LCD). Notable examples are FUS, TPD43, α-synuclein and nhRNPA1. All of these readily undergo liquid-liquid phase separation^[5]^ and in the early maturation stage reversibly interact through short, non-amyloid β-sequence motifs, from which irreversible amyloid aggregates can develop.^[6], [7]^ The fact that the latter are believed to be neurodegenerative^[8]^ makes physically understanding even pre-amyloid ordering or non-amyloid aging of direct pathological interest.

Although not a classical neurodegenerative disease-associated protein, the IDP Nucleoporin-98 (Nup98) has been implicated in neurogenerative diseases^[9]^ and can, like most IDPs, transform to amyloid states.^[10], [11], [12]^ Like many FG-rich Nups, this protein has been found to be capable of phase separating to yield a dense phase, which is typically initially liquid but becoming more viscous over time.^[13]^ A major feature of the NPC’s permeability barrier is that it excludes passage of cargoes, the bigger they are, unless they can bind to transport receptors like Importin-β, which facilitate interaction with the barrier and can guide transport through the pore. The nature of the *in-vivo* barrier is still under debate, as is the functional need for hydrogel formation versus the liquid state. However, both the liquid and gel states are good *in vitro* model systems for the actual barrier function of the full NPC.

IDP phase separation has been successfully modeled using the *associating polymer theory* introduced by Semenov and Rubinstein nearly three decades ago.^[14], [15]^ This free energy-based approach describes an interacting heteropolymer in terms of ‘stickers’, *i.e*. subunits that associate via a specific binding motif and ‘spacers’ which comprise monomers that interact with their environment in a non-specific manner. Since the establishment of the biocondensate field, extensions of the theory have appeared, including multiple sticker types,^[16]^ long range electrostatic interactions,^[17]^ crowding and higher order monovalent association,^[18]^ or an additional associating polymeric component, in the melt^[19], [20]^ or in solution.^[21], [22], [20], [23]^ The theory is attractive as it provides an intuitive thermodynamic picture of phase behavior, percolation and sol-gel transition, at least qualitatively reproducing virtually all currently documented biopolymer phase diagrams and often compares favorably with simulations.^[23]^ In a recent extension of the theory,^[24]^ the effect of ATP on the phase separation of fused in sarcoma (FUS) has been investigated, placing the specific interaction between Arg and Tyr residues in competition with binding of ATP to the former via an adaptive binding stoichiometry. As a result, re-entrant phase behavior could be described as a function of the ATP concentration.

A shortcoming of associating polymer-based models, is that the sticker valency is fixed rather than subject to how an IDP naturally interacts with its environment. These models therefore fail to capture an IDP’s adaptive molecular properties, such as (templated) folding,^[25], [26]^ making them unsuitable to computationally study maturation and aging. Simulations have shown that the presence of strongly interacting stickers can lead to a dynamic arrest^[27], [28], [29]^ or solidification,^[30]^ but do not model the increase in the stickiness of the IDP chains induced by conformational changes. For the same reason, associating polymer theory-based models, though effective in predicting IDP phase and percolation behavior of *native* biocondensates, cannot describe maturation and aging. Presently no intuitive theoretical framework exists that interrelates the driving forces for phase separation and aging in a unified manner.

This work aims to amend this by presenting a thermodynamically consistent model for *in-vitro* phase separation and β-sheet-type aging based on a *time-dependent, multi-component extension of associating polymer theory* in which binary associating sites form in time as a function of protein solubility, association strength, binding partners and folding propensity. As such, our coarse-grained model describes the aging process in terms of an increase in time of the *valency* of strongly interacting binding sites. This choice is based on the notion that an increase in valency i) within our model represents (the consequence of) conformation ordering associated with β-folding and ii) leads to an increase in the effective binding strength, as well as the contact life time, both between single stickers and between chains. We provide an intuitive explanation for the spontaneous increase in the ‘stickiness’ of an IDP and concomitant rise in viscoelasticity. Owing to its inclusive thermodynamic description, not only demonstrates in what way phase separation affects aging kinetics, but also how interactions forming during aging can induce phase separation. The model is consistent with well-known bio-recognition motifs including conformational selection, induced fit,^[25]^ and multi-valent fuzzy association^[31]^ and complies with what has recently been dubbed “COAST”, *i.e*. the ability of biopolymers to exhibit *coupled associative and segregative phase transitions*.^[32]^

We demonstrate the biological relevance of our model by validating our calculations against IDP aging essays based on the “perfect repeat” version of Nucleoporin-98 (Nup98),^[33]^ a model system originally designed to maintain the functional capabilities of Nup98 to mimic NPC behavior with minimal sequence complexity. Namely, 52 repeats of the 12 amino acid sequence GGLFGGNTQPAT, containing a key FG residue was found to capture the key NPC like permeability properties of wild type Nup98 droplets. As specific mutations like the mutation to V in the inter FG region have been found to promote gel formation, we hypothesize that these drive aging by contributing to the formation of β-sheets that act as a stickers that can form over time. This makes the perfect repeat variants of Nup98 excellent model systems to study and tune the impact of stickers during aging of any IDP that ages via folding-induced β-sheet based amyloid or non-amyloid interactions.

The remainder of this paper is set up as follows: in Sections 2.1 and 2.2 we detail the model and present two scenarios for how phase separation and aging are coupled. Phase separation encourages aging through increased local concentration but may, depending on the properties of the IDP, itself be driven by the aging process. In Section 2.3 we demonstrate thermodynamic consistency and delineate how the aging kinetics is affected by the molecular properties and solubility of an IDP. Our calculations predict a strongly non-linear dependence of the aging kinetics on sticker valency. In Section 2.4 we validate our model against experimental aging of V-mutants of perfect repeat Nup98-based IDP condensates, showing semiquantitative agreement between measured and calculated aging dynamics. In Section 2.5 we calculate the viscoelasticity of (aged) droplets and compare our calculations with dynamic data from literature. We predict viscosities of the order 10 to 100 Pa·s and pronounced viscoelastic behavior even for only moderate sticker association strength and without having to assume the formation of extensive fibrous networks. An extended summary and some additional discussion are given in Section 3.

## 2. Results and Discussion

### 2.1. Theory and Model

As mentioned in the Introduction, within our model aging is represented by the time-dependent conversion of precursor sites into sticker sites, which increases the average sticker valency with time and gives rise to an effective multicomponent solution, comprising polymer species with different valencies, to which we will refer as the *valency distribution*. Our model considers a solution of a disordered protein bearing *ξ*_*n*_ precursor- or sub-sequences along its backbone (see Figure 1A), of which each can, via a local and reversible conformational change or fold, switch between a ‘disordered’ precursor and a locally ‘ordered’ sticky site, that we will refer as a ‘sticker’. Each sticker is capable of forming a binary association complex with another sticker. The number of precursor sequences per chain is not principally limited, but constrained to be much smaller than the total number of monomers *N* per chain, so *ξ*_*n*_ << *N*. The precursor sites are interspersed by spacers comprising monomers that (co-)determine the phase behavior of the IDP via non-specific interactions, such as π-π, cation-π, aromatic or hydrophobic interactions. Although in some literature, these residues would themselves be identified as ‘stickers’, in this work we deliberately reserve this term for said associating sites, consistent with the terminology of Semenov and Rubinstein. We absorb the typically weak, non-specific interactions in binary interaction parameters (*χ*_*ij*_), consistent with classical models for the phase behavior of polymer mixtures and solutions, such as Flory-Huggins theory.^[34]^

**Figure 1.**
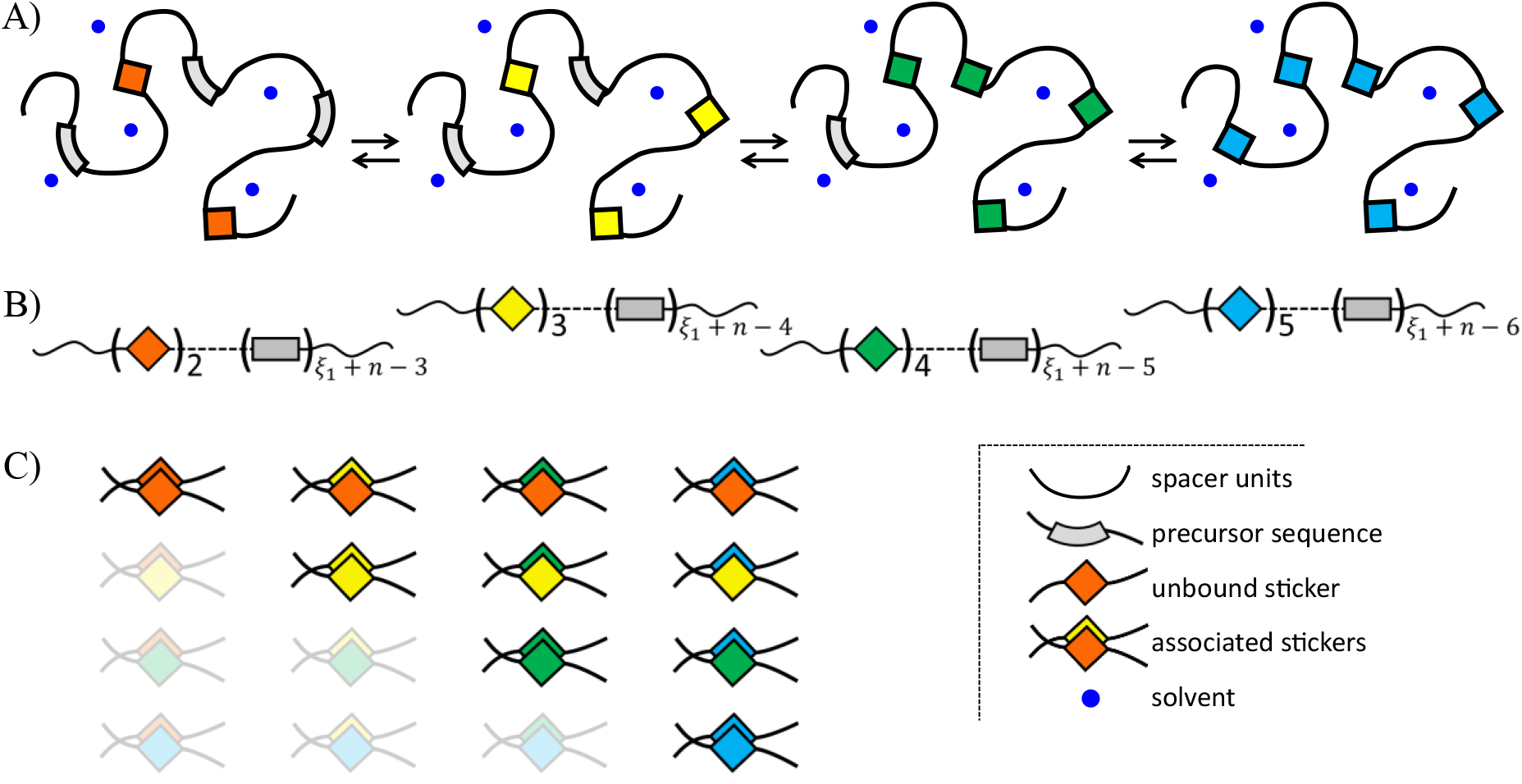
Schematic representation of A) *n* = 4 different but interconvertible IDP species (chain types) with a minimum and maximum valency of and *ξ*_1_ = 2 and *ξ*_*n*_ = 5 stickers. B) Shorthand representation of each species. The grey rectangles represent precursor units and the index relates the number of precursor units to the minimum active sticker valency *ξ*_1_ and the total number of different IDP species *n*. C) Symmetric matrix showing the number of different pairwise sticker complexes (multiplicity) that can form between the polymers shown in A). The number of different binding equilibria does not depend on valency but does depend on the number of species as *n*(*n* + 1)⁄2.

The interconversion between precursor sites and stickers modulates the number of different species in solution, which has to be accounted for entropically. In other words, despite the fact that we consider a single IDP, the solution becomes a multicomponent mixture containing *n* different species, each having a different sticker valency *ξ*_*i*_ in the range *ξ*_1_ ≤ *ξ*_*i*_ ≤ *ξ*_*n*_, with *ξ*_1_ ≥ 2 being the minimum valency. So, *ξ*_*i*_ = *ξ*_1_ + *i* − 1 and *ξ*_*n*_ = *ξ*_1_ + *n* − 1. As any free energy based on the theory by Semenov and Rubinstein,^[14]^ our thermodynamic model neglects sequence specificity and treats stickers as uncorrelated. As shown in numerous earlier studies,^[16], [17], [18], [19], [20], [21], [22], [23]^ this does not in any way limit its applicability to identifying and elucidating the thermodynamic drivers behind phase- and constitutive transitions characteristically observed for bio- and heteropolymers. Below we demonstrate and validate that this approach forms an ideal starting point, not only for constructing a unifying thermodynamic framework that includes aging, but also for modelling changes in constitutive properties, where sequence specificity is in fact taken into account. Figures 1A and B schematically depicts an exemplary system with *n* = 4 and 2 ≤ *ξ*_*i*_ ≤ 5, the B-panel presenting a ‘short-hand’ representation, which we clarify in the accompanying caption. The spacer units contain the non-specifically interacting monomers, which we do not explicitly show in the graphical representation.

We describe the above system in terms of a free energy as a generalization of the associating polymer model for a single species with a fixed number of specifically interacting stickers, originally proposed by Semenov and Rubinstein^[14]^. The following simplifications and assumptions apply: i) no correlations between stickers by demanding that they are spaced relatively far apart, ii) no consideration of chain connectivity and iii) the total polymer concentration exceeds the overlap concentration (semi-dilute regime). Under these conditions, we write the dimensionless mean-field free energy for the (*n* + 1)-component mixture as:

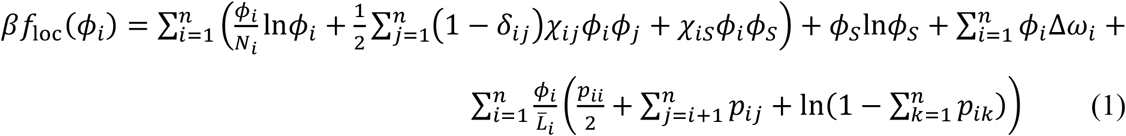

with *β* = 1⁄*k*_*B*_*T* the inverse thermal energy and *k*_*B*_ Boltzmann’s constant. The subscript “loc” refers to the fact that the above free energy applies to a homogeneous system, determined by the polymer volume fractions *ϕ*_*i*_ as local quantities.

The first two terms on the RHS of Equation 1 represent a multicomponent Flory-Huggins mixing free energy with *ϕ*_*S*_ ≡ *ϕ*_*n*+1_ the volume fraction of solvent or buffer. The mixture is treated incompressible, so that *ϕ*_*S*_ = 1 − *Φ*, with 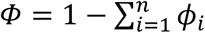 the total polymer volume fraction. *N*_*i*_ denotes the size of species *i*, normalized to that of the “solvent”: *N*_*S*_ = 1. *δ*_*ij*_ is the Kronecker delta. *χ*_*ij*_ and *χ*_*iS*_ represent the interaction of the regular monomers with their surroundings, the former representing non-specific interactions between polymer species and the latter being the polymer-solvent interaction parameter or ‘solvency’. In some calculations the latter is formulated as: *χ*_*iS*_ = *A* + *B*⁄*T*, with *A* and *B* empirical constants,^[35]^ typically evaluated based on experimental data.

The third term on the RHS represents a contribution associated with the local conformational change required to transform a precursor site into a sticker. This contribution is given relative to that of the lowest valency state: Δ*ω*_*i*_ = (*i* − 1)Δ*ω*_0_, with Δ*ω*_0_ the energy change for the bare conformational transition that gives rise to an active sticker. Although when projected onto β-folding, Δ*ω*_0_ is strictly a sum of energetic and entropic contributions, we do not specify these separately. Instead, we treat this quantity as a control parameter and assume that the chain gains free energy upon sticker formation, so Δ*ω*_0_ > 0. This choice gives rise to an interesting competition between an unfavorable conformational contribution, a favorable translational entropic contribution (creating as many different species as possible) and a favorable energetic contribution due to sticker association. So, by convention: i) the degree of conformational ordering of a polymer increases with the number of active stickers and ii) in the absence of sticker binding the largest energetic cost is associated with the formation of the species with the highest valency.

The last term in Equation 1 is the generalization of the model by Semenov, Rubinstein and others^[14], [19], [20], [21], [22]^ to a mixture or solution of any number of species comprising any number of pairwise associating stickers, spaced along the backbone by an average number of 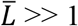 monomers. Its derivation is given in Section S1 of the Supporting Information (SI) The ratio 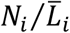 simply represents a proportionality factor that expresses the valency of the *i*^th^ polymer species. The parameters *p*_*ij*_ denote the equilibrium fractions of stickers of the *i*^th^ chain type involved in a pairwise complex with a sticker of the *j*^th^ chain type. Semenov and Rubinstein showed that minimizing a free energy based on the sum of a translational entropy of the sticker-free reference chains and a contribution due to sticker binding multiplicity, such as Equation 1, with respect to the (independent) bound fractions *p*_*ij*_ is equivalent to minimizing a free energy that includes the translational entropy of all association clusters with respect to the cluster size, as long as these are assumed tree-like and the minimum sticker valency *ξ* ≥ 2.^[14], [20]^ Furthermore, since we (mostly) consider a concentrated droplet state, in which excluded volume interactions are screened and chains behave virtually ideal, a reduction in the contact probability between stickers due to chain swelling is not considered.

The fractions of stickers in a given bound state are related to the molar association constant *K*_*ij*_ according to: 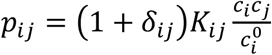, with *c*_*ij*_ the molar concentration of free stickers of species {*i, j*} in the binding equilibrium, 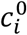 the total molar concentration of stickers of species *i*. The association constant is related to a binding energy *ε*_*ij*_ as: *K*_*ij*_ = *v*_*m*_exp(*βε*_*ij*_), with *ε*_*ij*_ ≥ 0 and *v*_*m*_ a reference molar volume of *v*_*m*_ = 1 l/mol, producing the correct dimensions as *K*_*ij*_ is expressed in l/mol. We relate 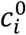 to the volume fraction of *i*-polymers as: 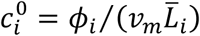. Dividing *v*_*m*_ by Avogadro’s number 𝒩_*A*_ yields a ‘monomeric’ volume *v*_0_ = *v*_*m*_⁄𝒩_*A*_ which forms the basis for the normalization of molecular size. Up to a prefactor of order unity we obtain a suitable fundamental length scale *b* = (*v*_*m*_⁄𝒩_*A*_)^1⁄3^ ≈ 1.1 nm, which coincides very well with the Kuhn length of a flexible polymer, such as an IDP. The molar concentration of chains of type *i* is related to the volume fraction as: *C*_*i*_ = *ϕ*_*i*_⁄(*v*_*m*_*N*_*i*_), with *C*_tot_ = ∑_*i*_ *C*_*i*_ the total molar polymer concentration. The details on the calculation of all concentrations and association probabilities within the multicomponent binding equilibrium are given in Section S2 of the SI.

The temperature dependence of the association constant is incorporated by parametrizing the binding energy in terms of an energetic and an entropic contribution according to:

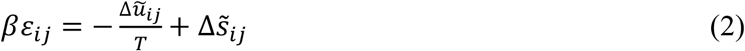

with 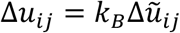 and 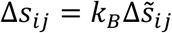 the energy and entropy of sticker binding. For clarity, 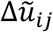 is a dimensionless quantity and 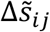 is given in units of absolute temperature. The binding energy co-determines the time a sticker of a given chain resides in the associated state. The bare life time of an *ij*-sticker complex is given by its dissociation time^[15]^ 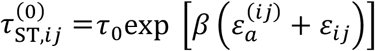, with 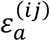 the energy barrier for association, *τ*_0_ = *b*^2^⁄*D*_0_ and *D*_0_ the relaxation time and self-diffusivity of a regular monomer or solvent molecule.

During an *in-vitro* experiment involving phase separation and aging no energy is consumed or generated. In other words, condensate formation and subsequent aging are passive processes and should principally be accompanied by a perpetual decrease in free energy towards a minimum, at which point detailed balance must be obeyed.^[36], [37]^ This implies that the driving force for sticker formation is provided by the exchange chemical potential 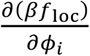.^[38]^ Following Weber *et al*.^[39]^ and assuming the local folding and unfolding events required for generating and resolving stickers to be elementary unimolecular reactions, we may write the reaction fluxes for the formation of the different species according to a Becker-Döring-type process: a chain with *ξ* stickers forms via a folding event in a chain with *ξ* − 1 stickers or an unfolding event in a chain containing *ξ* + 1 stickers, such that:^[40]^

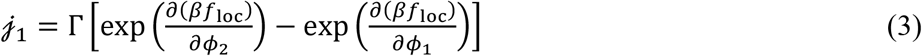

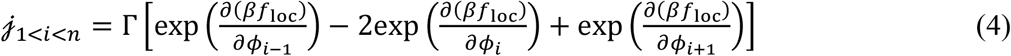

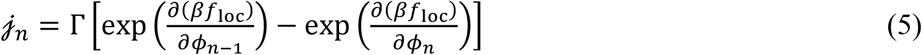

The (Onsager) reaction mobility Γ may be formulated based on an effective Arrhenius relation 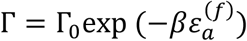, where the value of 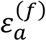 represents the barrier for the equilibrium exchange between local folding and unfolding. The reaction mobility is divided out in our calculations, as we express aging as a function of dimensionless time 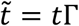. It only becomes of importance where we treat dynamic arrest, where the effective activation barrier diverges (Section S8 of the SI).

On account of the fact that the precursor segments are much shorter than the length of a strand between them (*ξ*_*n*_ << *N*), we neglect changes in molecular volume upon sticker formation, which means *N* is conserved. As we will show in Section 2.3, the fluxes in Equations 3-5 evolve towards an equilibrium, wherein the total free energy in Equation 1 is minimized and all chemical potentials equalize and no net flux occurs. At this point, an equilibrium distribution of species with different sticker valencies is reached, which to which we refer as the *equilibrium valency distribution*. It is noted that this distribution may be subject to thermal fluctuations, which are not considered by the present model due to its mean-field nature. If dynamic arrest is considered (see Section S8 of the SI), Equations 3-5 lead to an out-of-equilibrium state where the net flux is negligibly small, though now due to kinetic frustration.

The fact that the fluxes of active stickers and precursors depend on the exchange chemical potentials 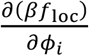, implies that the associated reversible conformational transitions are influenced by everything that contributes to the free energy. In other words, besides the ‘bare’ energy change associated with the conformational transition, also concentration, chain length, translational entropy, solvent interaction, binding strength and binding probability all together determine the driving forces for local folding and unfolding. All input parameters for the calculations presented in the remainder of the work are listed and described in Tables S2 – S4 in Section S6 of the SI. Besides, to support the presentation and discussion of the results, we repeat and specific parameters in the main text and figure captions.

### 2.2. Phase Behavior

In this section we study how ‘aging’, *i.e*. the formation and association of stickers on an IDP in solution, influences the binary temperature-composition phase diagram. To assess what the implications are of a particular biomolecular phase behavior and to demonstrate the model’s mechanistic scope and applicability, we consider two scenarios. In Scenario I, phase separation of the IDP is predominantly driven by a strongly unfavorable solvency (embodied by a high *χ*_*iS*_, and to much lesser extent by the polymer becoming ‘stickier’ due to aging, whereas in Scenario II we assume the solvency rather favorable, *i.e*. lower *χ*_*iS*_ with sticker formation and association *becoming* the primary driving force for phase separation. We will show that our model handles both scenarios naturally and that both scenarios are physically and biologically relevant.

Scenario I, schematically depicted in Figure 2A, mimics a typical *in-vitro* phase separation experiment in which IDP-rich droplets form relatively fast, for instance in response to a buffer exchange^[41]^ or an enzymatic cleavage of a solubilizing tag, such as maltose binding protein (MBP).^[42]^ At the moment of droplet formation we consider the IDP to be in its unaged, most disordered state, here: lowest valency state, indicated with orange stickers (middle panel of Figure 2A). Subsequently, the IDP slowly ages (mostly) in the condensed phase, through the spontaneous development of higher valency species (right panel). The timescales of phase separation and aging are vastly different and the two processes occur *sequentially*. Nevertheless, aging is accelerated by the increase in local concentration. Owing to the separation of time scales of phase separation and aging, Scenario I allows for modeling condensate aging without having to (explicitly) consider mass transport, under the assumption that the IDP concentration in the condensates does not change over time, which we will show to be the case.

**Figure 2.**
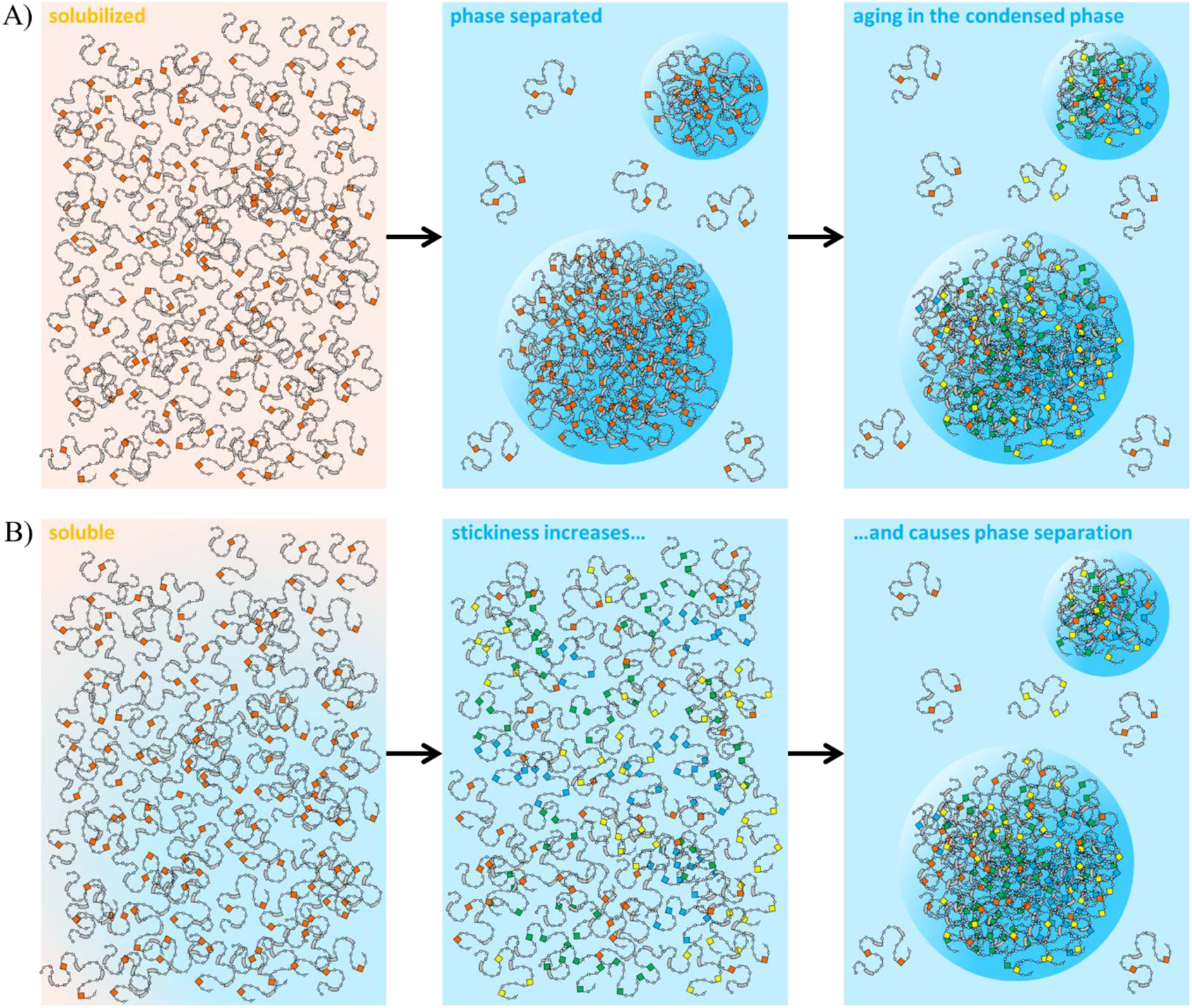
Schematic representation for two scenarios for phase separation and aging, both compatible with our model. A) Scenario I: IDP phase separation is driven by an induced poor solvency (high *χ*_*iS*_) and B) Scenario II: phase separation is driven by aging itself. In A) phase separation of the initial (most denatured/disordered) species (here with *ξ*_1_ = 2, orange species) takes place, for instance following a buffer exchange or enzymatic cleavage of a solubility tag. Subsequently, spontaneous aging occurs in, predominantly, the concentrated phase. In B) the aging increases the stickiness of the still soluble IDP, at some point in time leading to phase separation. Formation of higher valency, ‘aged’, material is in both panels indicated in yellow (*ξ*_2_ = 3), green (*ξ*_3_ = 4) and blue (*ξ*_4_ = 5).

In Scenario, II (Figure 2B) the IDP remains well-solvated as long as sticker formation, *i.e*. ‘aging’, has not progressed substantially (left and middle panel). In this scenario, aging *may* also be slower than phase separation but, more importantly, the latter occurs *as a consequence of the former* (right panel). Scenario II is perhaps more representative for systems where sticker-sticker interactions are responsible for both phase separation and the increase in the viscosity on account of slow conformational changes that maximize sticker interactions.^[43]^ In contrast to Scenario I, Scenario II can only be properly modeled by taking mass transport explicitly into account, which we will address in future work.

The way in which Scenarios I and II differ is illustrated by the model’s prediction of how the dense phase compositions *C*_con_ is expected to change during aging. To inspect this, we, for now, consider a simplified case, where aging is coarsely represented by a stepwise increase in a fixed valency, each represented by a separate phase diagram (Figure 3). This exercise does *not* give us the coexisting equilibrium *valency distributions, i.e*. the concentration of the different species within each phase, which we address separately below. The phase diagrams in Figure 3 are calculated using a binary common tangent construction based on Equation 1, for a given valency (2 ≤ *ξ* ≤ 8, chain length *N*, association strength *K* and solvencies (*χ*_*iS*_) (all input is given in Section S6 of the SI). The common tangent construction equalizes the exchange chemical potential ∂*f*_loc_⁄∂*ϕ* and osmotic pressure *ϕ* ∂*f*_loc_⁄∂*ϕ* − *f*_loc_ in both phases. To demonstrate the generality of our argument, we considered both upper critical solution temperature behavior (UCST) and lower critical solution temperature behavior (LCST). LCST behavior is less commonly encountered in biological phase transitions than UCST, but has for instance been observed for FG-Nups^[44]^ and elastin-like polypeptides.^[45]^

**Figure 3.**
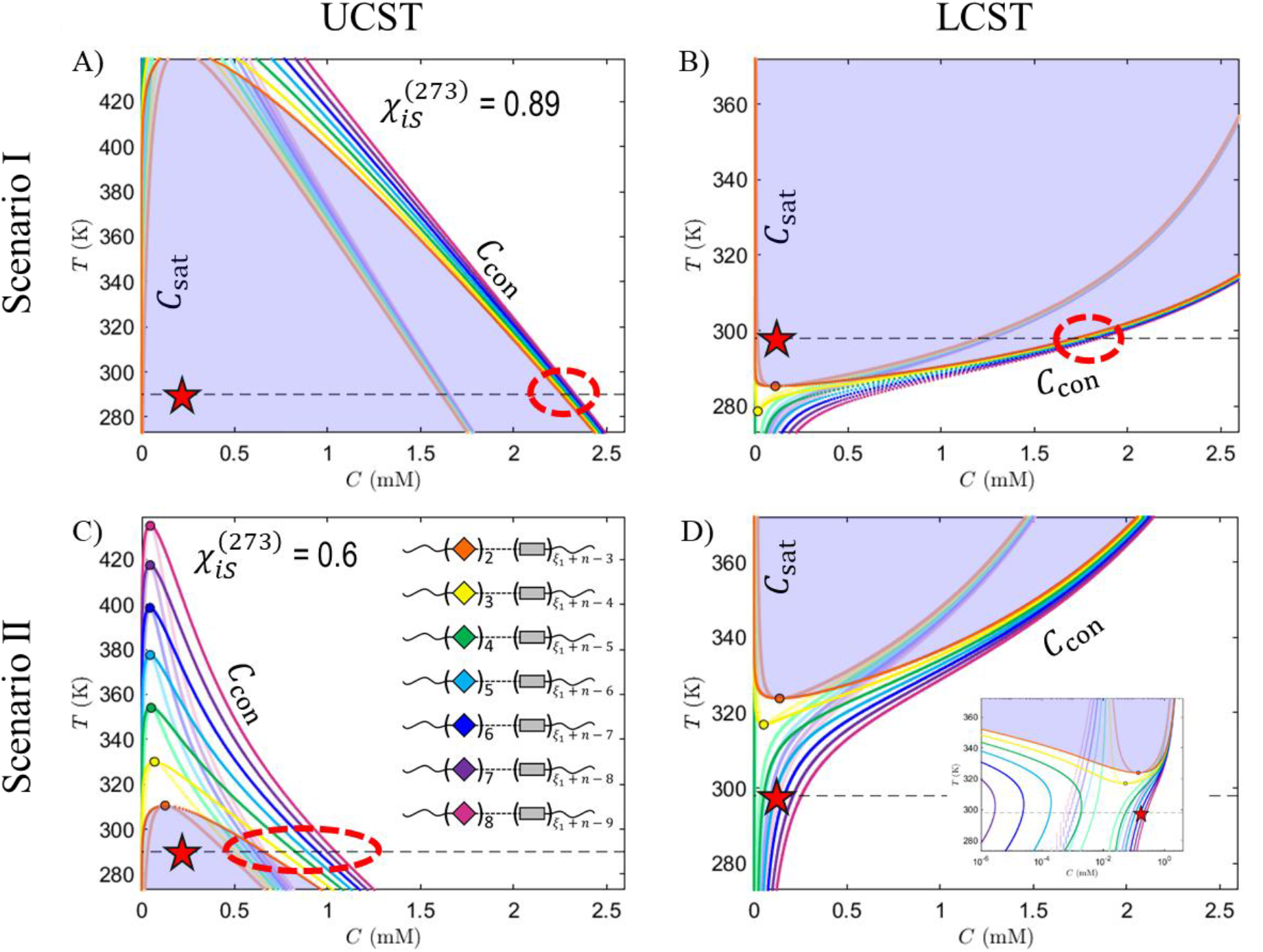
Temperature-composition phase diagrams, calculated using Equation 1 for a fixed valency ranging from bivalent (*ξ* = 2) up to octavalent (*ξ* = 8, *n* = 7) disordered proteins (legend in C) with panel pairs A, B and C, D respectively representing Scenarios I and II (see main text) for UCST (A and C) and LCST (B and D) behavior. The symbols, dark lines and shaded lines respectively represent the position of the critical point, binodal and spinodal for each valency. The parametrization is as follows: *N* = 250,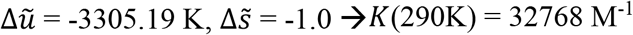. *χ*_*iS*_(*T*) = 0.89 × 273⁄*T* (panel A), 0.6 × 273⁄*T* (panel C), 5.0397 − 1288.5/*T* (panel B) and 2.5198 − 644.24/*T* (panel D). The dashed black lines indicate temperatures of 290K (A and C) and 298K (B and D). The red stars are on the dashed lines at an arbitrary mean IDP concentration. The inset in panel D represents the same phase diagrams, plotted semi-logarithmically.

The temperature-dependence of both the non-specific interactions, encoded by *χ*_*iS*_(*T*) = *A* + *B*⁄*T* (see above, and the specific interactions encoded by *K*(*T*) = *v*_*m*_exp(*βε*_*ij*_), allows for these interactions *by themselves* to already produce UCST or LCST behavior. The expression for *χ*_*iS*_(*T*) produces UCST for *A* > 0 and *B* > 0 and LCST for *A* > 0 and *B* < 0. In case of the specific interactions UCST and LCST can, per Equation 2, be respectively obtained by energy-driven complexation (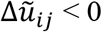 and 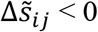) or entropy-driven complexation (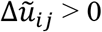 and 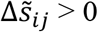). As our model combines non-specific and specific interactions, it is capable of producing a very rich phase behavior based on four possible permutations, which we partly exploit in Figure 3. We have (non-specific/specific) UCST/UCST in Figure 3A and 3C, and LCST/UCST in Figure 3B and 3D. Hence, Figure 3 assumes LCST behavior to stem from non-specific interactions and does not show cases of phase separation by entropy driven sticker-sticker binding, simply because it is not within its scope and bypasses its intention of demonstrating the different aging scenarios. We assign biologically relevant input by assuring that the relevant phase behavior coincides with a physiological temperature (see Section S6 of the SI).

The dark colored lines in each panel demarcate the coexistence region and represent the (binodal) concentrations of the dilute (*C*_sat_) and condensate phase (*C*_con_), that coexist for a given temperature. The lines in corresponding lighter hue, obtained by solving the condition ∂_*ϕϕ*_(*βf*_loc_) = 0, represent the spinodal curves, demarcating the region of thermodynamic stability. The full parametrization is given in the caption. The red star in each panel represents an exemplary initial condition with the IDP in its unaged, lowest valency state (*ξ* = 2), *i.e*. right after buffer exchange or tag cleavage (Scenario I, panels A and B) or simply in solution (Scenario II, panels C and D). Depending on the solvency, the initial condition is either relatively deep inside the coexistence region (panels A and B), near the critical point (panel C) or outside the coexistence region (panel D). If it is inside the coexistence region (*i.e*. panels A-C), the solution phase separates along the black dashed tie-line to reach the binodal at *C*_sat_ and *C*_con_. However, since, irrespective of the Scenario, aging sets in as well, the IDP becomes stickier and hence less soluble, which widens the coexistence-region (see binodals of subsequent color). Consequently, the red star is located ever deeper inside the miscibility gap and further away from the critical point.

In Scenario I, this expansion of the coexistence region has a negligible effect on *C*_con_ (panels A and B, dashed ellipses), for which reason the phase diagrams in Figure 3A and 3B would look very similar if we would not have fixed the valency at a single value, but instead have treated it as a natural function of the total concentration. In other words, *C*_con_ is determined by molecular size and solvency, not sticker valency. Hence, Scenario I allows for modeling condensate aging without having to (explicitly) prior phase separation and by approximation assuming the total dense phase concentration to remain constant over time. We will make use of this in subsequent calculations, for which a constant IDP volume fraction is given by the binodal of the relevant lowest valency species (see Table S2 in Section S6 of the SI).

In Scenario II, a shift in the dense phase concentration is considerable. *C*_con_ becomes a ‘moving target’ (Panel C) and only well-defined when an equilibrium interface appears between the two coexisting phases, at which point in fact a Scenario I would have emerged. In Panel D phase separation only sets in if the (average) valency of the IDP has increased up to the point that the star is situated inside the coexistence region (here reached for *ξ* > 5). In case of LCST behavior (panels B and D), *χ*_*iS*_ and *K* have an opposite temperature dependence, due to which the phase diagram becomes reentrant for valencies *ξ* > 3. This is apparent if we plot the phase diagram in Figure 3D on a semilogarithmic scale (inset). We emphasize that, in contrast to Scenario I, although the representation in terms of phase diagrams based on a stepwise increase of a single valency effectively shows the shift in dense phase composition, the profound non-equilibrium nature of the early stages of Scenario II is actually hard to capture in (a series of) phase diagrams. Interestingly, Scenario II is somewhat reminiscent of a volatile solution that destabilizes due to evaporation of one of the components, driving the composition deeper into the miscibility gap in a so-called ‘imperfect quench’.^[46], [47], [48]^

Once global equilibrium has been reached, whether in Scenario I or II, we can determine the valency distribution in the coexisting phases by applying the common tangent construction to the full free energy Equation 1, while treating the equilibrium valency distribution as a natural function of the total polymer concentration, rather than fixing a specific single valency. These calculations, of which the results are shown in Figure 4, superimpose two additional layers of complexity compared to the calculations underlying Figure 3: i) solving the full *multicomponent binding equilibrium* to evaluate the bound sticker factions (see Section S2 of the SI) and ii) obtaining the equilibrium valency distribution as a function of total concentration. The latter is obtained numerically integrating Equations 3-5 until all fluxes become zero. The actual common tangent construction is applied to a polynomial fit to the free energy, of which we show the first derivative with respect to concentration, overlayed to the total exchange chemical potential ∂*f*_loc_⁄∂*Φ*.

**Figure 4.**
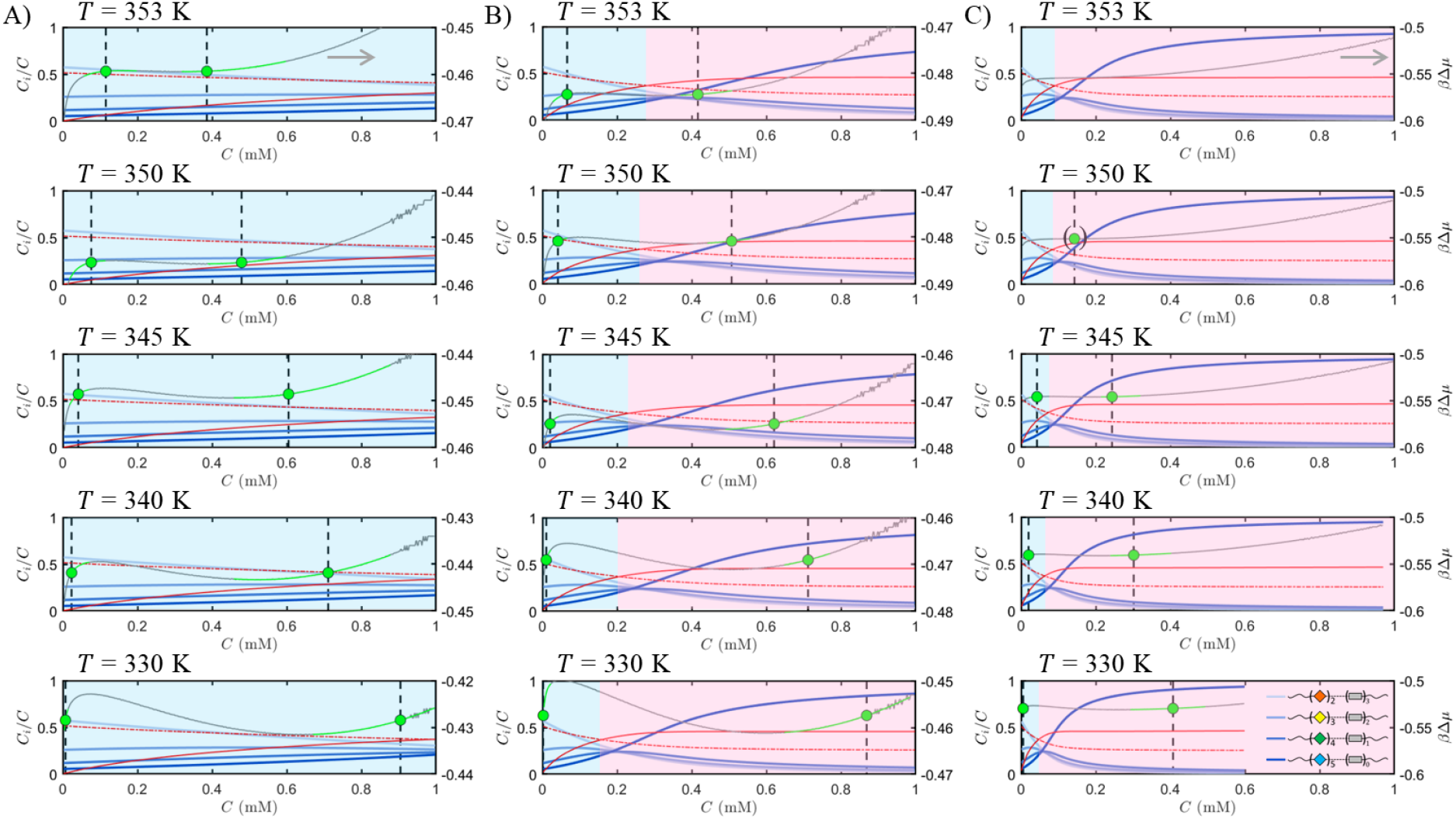
Equilibrium binodal compositions as a function of temperature, binding strength and solvency. In A) phase separation is predominantly driven by a poor solvency; in C) phase separation is predominantly driven by sticker association; in B) binding and solvency contribute about equally. The blue curves represent the equilibrium concentration of the IDP species *C*_*i*_ with different sticker valencies (see legend in C, 330 K), normalized by and plotted as a function of the total molar polymer concentration *C*. The green symbols and vertical dashed lines indicate (from left to right) the coexisting compositions *C*_sat_ and *C*_con_. In C) for *T* = 350K, they approximately indicate the critical point. The grey curves represent the exchange chemical potential. The ‘green sleeves’ represent the first derivatives of polynomial fit functions to the free energy, used to determine the binodal compositions. The solid and dash-dotted red curves represent the total fraction of bound stickers and the mean-field percolation threshold, respectively. The blue and red shaded boxes indicate the non-percolated and percolated regions. The parametrization is as follows: A) 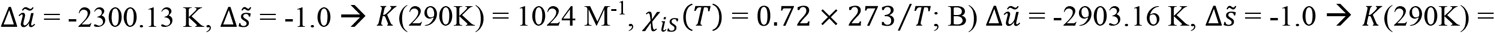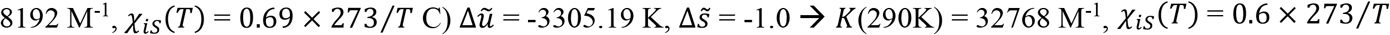; in all cases: *N* = 250 and *n* = 4.

In Figure 4 we consider three cases: one where phase separation occurs due to a poor solvency, while assuming weak sticker binding (Figure 4A), one where phase separation is driven by strong sticker association and less so by a poor solvency (Figure 4C) and one where both contribute about equally (Figure 4B). In the considered temperature range, the first case gives rise to almost similar coexisting valency distributions, whereas in the latter two the distributions are very different. In Figures 4B and 4C, the distributions are inverted: in the dilute phase it is dominated by the lowest valency species, whereas in the dense phase the highest valency species is most abundant. In each panel of Figure 4, the mean-field percolation threshold for the bound fraction, which for the present multi-component case, is given by:^[49]^

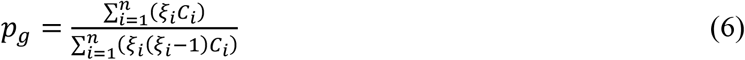

with *C*_*i*_ the concentration of IDP species *i* and *ξ*_*i*_ its valency. In the weak binding case (Figure 4A), percolation, indicated by the crossing of the solid and dash-dotted red lines, is not reached in either of the phases, whereas in the stronger binding cases (Figure 4B and 4C), the condensate phase comprises a dense percolated network, while the dilute phase remains non-percolated.

We end this section by providing some speculation on how aging would affect droplet coarsening in the two Scenarios sketched above and within the context of our model. Droplet coarsening can be due to i) coalescence, usually observed in the early stages of phase separation, and ii) Ostwald ripening, typically observed in late stages. In general, coarsening through coalescence can be hydrodynamic, giving rise to a linear time scaling of the mean droplet size as 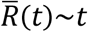.^[50]^ It may also occur diffusively, through Brownian collisions, which gives rise to a cubic root relation 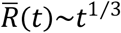 in 3D^[51]^ or 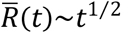 in 2D.^[52]^ Ostwald ripening is diffusive and exhibits the well-known Lifshitz-Slyozov-Wagner (LSW) scaling: 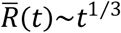.^[53], [54]^ Aging could potentially influence these processes, as the increased stickiness generally leads to a reduction in (diffusion) mobility and local viscosity,^[55]^ although only in the condensate phase.

For our Scenario I, wherein phase separation fast and based on a poor solvency, aging only becomes significant in the Ostwald ripening stage. We would not expect aging to influence ripening, since i) transport takes place through the (constant) dilute phase and ii) the LSW exponent follows from capillary and geometric arguments. Only if strong kinetic suppression leads to non-equilibrium concentrations near the droplets, it may appear the process loses its scaling universality. For Scenario II, sticker formation likely coincides with hydrodynamic mass-transport in the early stages of LLPS. Scaling behavior can be obscured due to various transient hydrodynamic regimes.^[55]^ Since aging would lead to an increased dynamic asymmetry, phase separation may even be viscoelastic, giving rise to an initial polymer-rich, network-like continuous phase.^[56], [57]^ This network breaks up due to capillary forces, upon which the dense phase becomes dispersed. Droplet coalescence might then occur, prior to late-stage Ostwald ripening. If coalescence is purely due to Brownian collision, we would expect little impact of aging as, again, mass transport takes place in the dilute phase.

### 2.3. Aging Kinetics

Next, we study aging of condensates that have just formed via phase separation of the lowest valency species according to Scenario I, *i.e*. for which phase separation is driven by a poor solvency and changes in *C*_con_ are negligible. As mentioned above, during aging a multicomponent mixture containing multiple IDP species with an increasing average valency develops in time. Hence, unless stated otherwise, all kinetic traces presented below start with a non-zero concentration of only the lowest valency species *ϕ*_1_ with generally *ξ*_1_ = 2. The initial concentrations of the other species are taken negligibly small (but non-zero to avoid divergence of logarithms). Furthermore, we note that since in the condensed phase the polymer concentration significantly exceeds its value at overlap, sticker association almost exclusively takes place on an inter-chain basis. Since all species in effect represent the same IDP, we assume the same binding strength for any pair of stickers, so *K*_*ij*_ = *K* and set the non-specific polymer-polymer interactions to zero: *χ*_*ij*_ = 0. Furthermore, in all calculations we fix the molecular size at *N* = 250 and the temperature *T* = 290 K.

The energy barrier for 1:1 sticker binding is assumed not to exceed a few times the thermal energy: 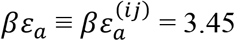. Against the applied ranges for *K* and the conformational energy penalty Δ*ω*_0_, values for Γ_0_ and 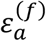 of, respectively, ~10^0^ s^−1^ and a few times *k*_*B*_ *T*, aging occurs on a much longer time scale than sticker binding and unbinding, which is a prerequisite for the validity of the associating polymer model, but breaks down under dynamic arrest as we discuss in Section S8 of the SI. The other input parameters for the calculations are detailed below, where we present a set of exemplary calculations that demonstrate how the composition and aggregation state of the IDP-rich droplet changes with time.

In our first example we inspect the effect of the solvency, expressed by *χ*_*iS*_. Figure 5 shows the aging of an IDP with a valency 2 ≤ *ξ*_*i*_ ≤ 5 (so *n* = 4) and a sticker association strength of *K* = 1024 M^−1^ for two solvencies: *χ*_*iS*_ = 0.68 (Figure 5A) and *χ*_*iS*_ = 1.13 (Figure 5B). Using Equation 1 we find that these values yield condensate concentrations of *C*_con_ = 1.49 mM and 3.00 mM for *ξ* = 2 and increasing by ~5% and ~0.4% for *ξ* = 5, respectively. The initial state (*t* = 0) represents condensates containing IDPs in their most denatured state, so *ξ*_1_ = 2. The blue traces in Figures 5A and 5B demonstrate how the valency distribution develops in time. Initially, the concentration of species with intermediate valency, *ξ*_2_ = 3 and *ξ*_3_ = 4 rise steeply at the expense of *ξ*_1_, and reach a maximum, after which the species with the highest valency, here *ξ*_4_ = *ξ*_*n*_ = 5, is eventually favored. Interestingly, the maxima in the traces for *ξ*_2_ and *ξ*_3_ are reached later in time if the solvency is worse. Importantly, Figures 5A and 5B show how, as a function of solvency, the average sticker valency of an IDP in a condensate *spontaneously* increases until an equilibrium is reached.

**Figure 5.**
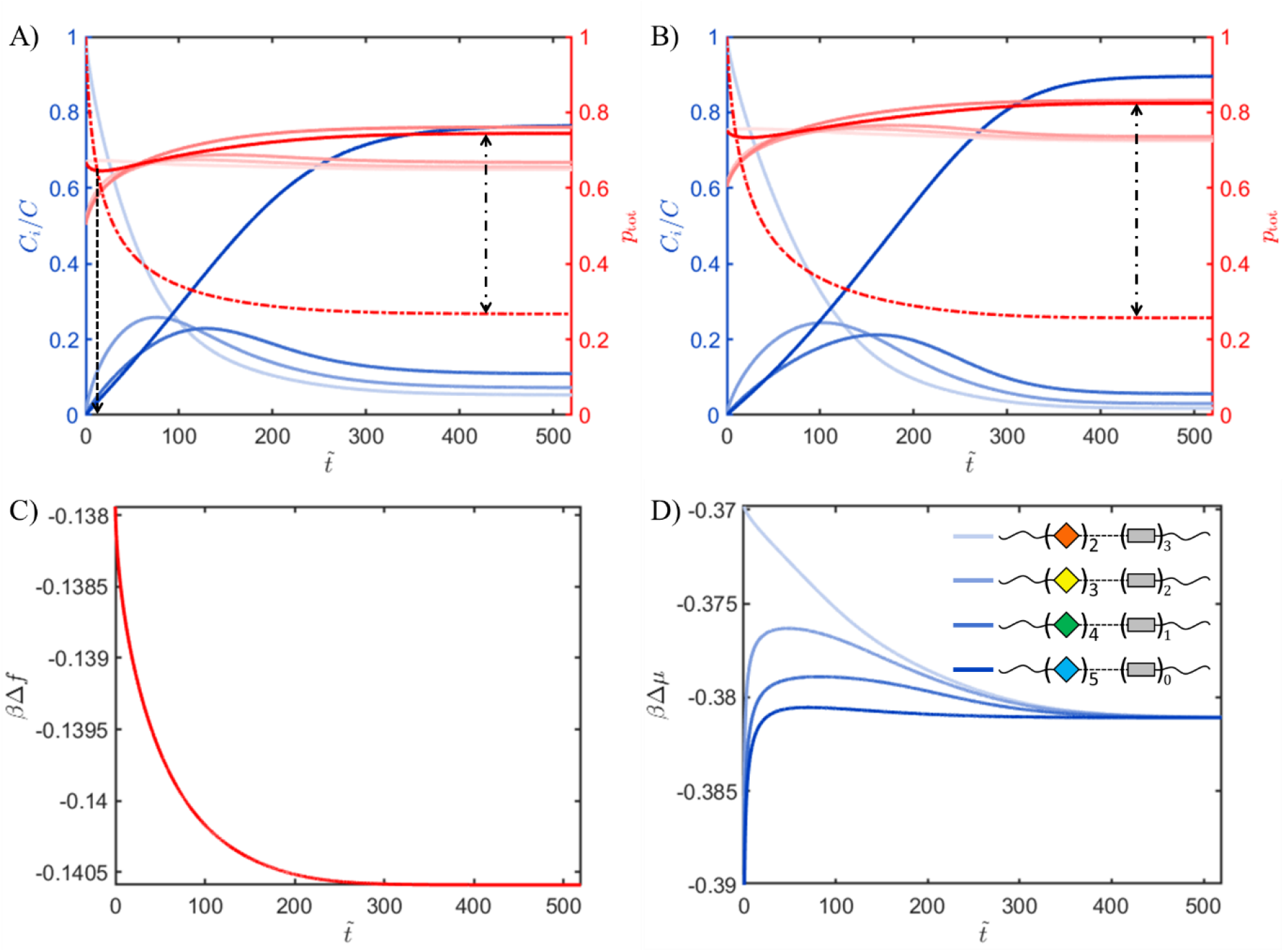
Kinetics and thermodynamics of aging in IDP-rich phase separated droplets. A) and B) concentration of IDP species *C*_*i*_ with different sticker valencies (see legend in D)), normalized by *C*_con_ (blue curves) and individual (red, shades) and total (red, dark) fractions of bound stickers; input for the calculations: *n* = 4, *ξ*_1_ = 2, *K* = 1024 M^−1^ and Δ*ω*_0_ = 0.0032, with *χ*_*iS*_ = 0.68 (A) and *χ*_*iS*_ = 1.13 (B), all plotted as a function of dimensionless time 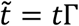. The dash-dotted red curves represent the mean-field percolation threshold and the dashed black arrows indicate the time to percolation. The dash-dotted black arrow expresses the difference between the total bound sticker fraction and its value at percolation. C) free energy per monomer in calculation A) plotted as a function of dimensionless time. D) chemical potential in calculation A) plotted as a function of dimensionless time.

Here, we purposely allow for aging to reach an equilibrium in order to study the thermodynamic driving force behind aging, but are aware of the fact that kinetic frustration may set in before equilibrium is reached. In Section 2.4 we provide some more discussion on dynamic arrest. Since the fraction of bound stickers and hence the free energy loss owing to association is determined by the product *K* × *C*_con_, the higher value for *χ*_*iS*_ yields condensates that are more strongly enriched in *ξ*_4_. So, more conformational order is expected at elevated concentrations, at which the binding equilibria are tending towards the bound state. Figure 5C and 5D demonstrate that the model is thermodynamically consistent: the free energy of the system decreases to a minimum upon reaching equilibrium and the chemical potentials equalize, meaning the net reaction fluxes, defined in Equation 3, 4 and 5, of all species become zero.

The solid red lines in Figures 5A and 5B represent the fractions of stickers in a bound state, calculated per individual species (red shades), as well as all species (*p*_tot_, dark red). Comparing Figures 5A and 5B shows that for lower solvent quality a larger fraction of stickers resides in a bound state, likely due to the higher condensate concentration. Therefore, the “stickiness” of the IDP not only depends on association strength *but also on concentration*. Our calculations show that *sticker-sticker association can drive the actual conformational switch required to form the stickers themselves*. The very nature of our model suggests that this does not necessarily require binding of the IDPs into extended (amyloid) fibers, a process we certainly do not claim to describe with our model: the phenomenon also applies to the formation of disordered, even reversible, aggregates or networks that are cross-linked via binary association between limited numbers of non-amyloid β-strands, as implicated in the development of neurodegenerative disease.^[7]^

Furthermore, a parallel can be drawn with the ‘local-nonlocal coupling principle’ in protein folding^[58], [59], [60], [61]^ in the sense that relaxation of the system relies on two elements: i) an interaction or binding between two distant/non-local sites and ii) a local conformational transition. The two processes depend on each other: the binding requires the conformational change and the conformational change can be driven by binding. So, irrespective of the sign of the energy change associated with the local conformational transition, it is the combination of the latter with a non-local interaction that forms the prerequisite that is to be fulfilled for the system to reach a lower energy state.

Irrespective of solvency or concentration, the total fraction of bound stickers *p*_tot_ (dark red curves) initially decreases and subsequently increases with time. This implies that if the concentration of higher valent species is very low (at early times), the density of unbound stickers initially increases faster than the density of bound stickers. As a result, in this regime the decrease in free energy is dominated by the gain in translational entropy associated with the population of multiple states in a broadening of the valency distribution, and not by complexation. At this early stage we would hence refrain from identifying the system as ‘aging’ since the bound sticker density, which is responsible for the constitutive properties of the solution, only marginally increases. The dashed black arrow in Figure 5A marks the point where the total bound fraction crosses the mean-field percolation threshold (dash-dotted red curve). At any time beyond indicated by the dashed black arrow, we have *p*_tot_ ≥ *p*_*g*_, giving a system-spanning network of associating chains, sometimes referred to as a ‘percolation transition’.^[32]^ As we show below, these calculations yield information that can be used to estimate the rheological properties of the network fluid in a regime where the high crosslink density is high. The latter is a function of the difference between the bound sticker fraction and the value at the percolation threshold (dash-dotted black arrows in Figure 5A and 5B).

Although assuming the lowest possible initial active sticker valence *ξ*_1_ = 2 is a reasonable choice from the perspective of biological aging, during which IDPs slowly transform into more folded and associated/bound structures,^[8]^ our model is not limited to describing this case and can in principle assume any concentration for any of the species within a valency distribution at 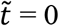. To demonstrate this, in Figure 6 we repeat the calculation performed in Figure 5A, but use a different abundant initial species in each panel: from the minimum *ξ*_1_ = 2 in Figure 6A up to the maximum valency component *ξ*_4_ = 5 in the distribution in Figure 6D. The same equilibrium valency distribution is reached with the relative concentration of each valency saturating at the same value, irrespective of the initial conditions. For an initial valency > 3, percolation is reached right from the start due to the suppressed percolation threshold (dash-dotted red curve). The total bound fraction (solid red line) flattens, meaning that for the highest initial valencies there is no net increase in *p*_tot_.

**Figure 6.**
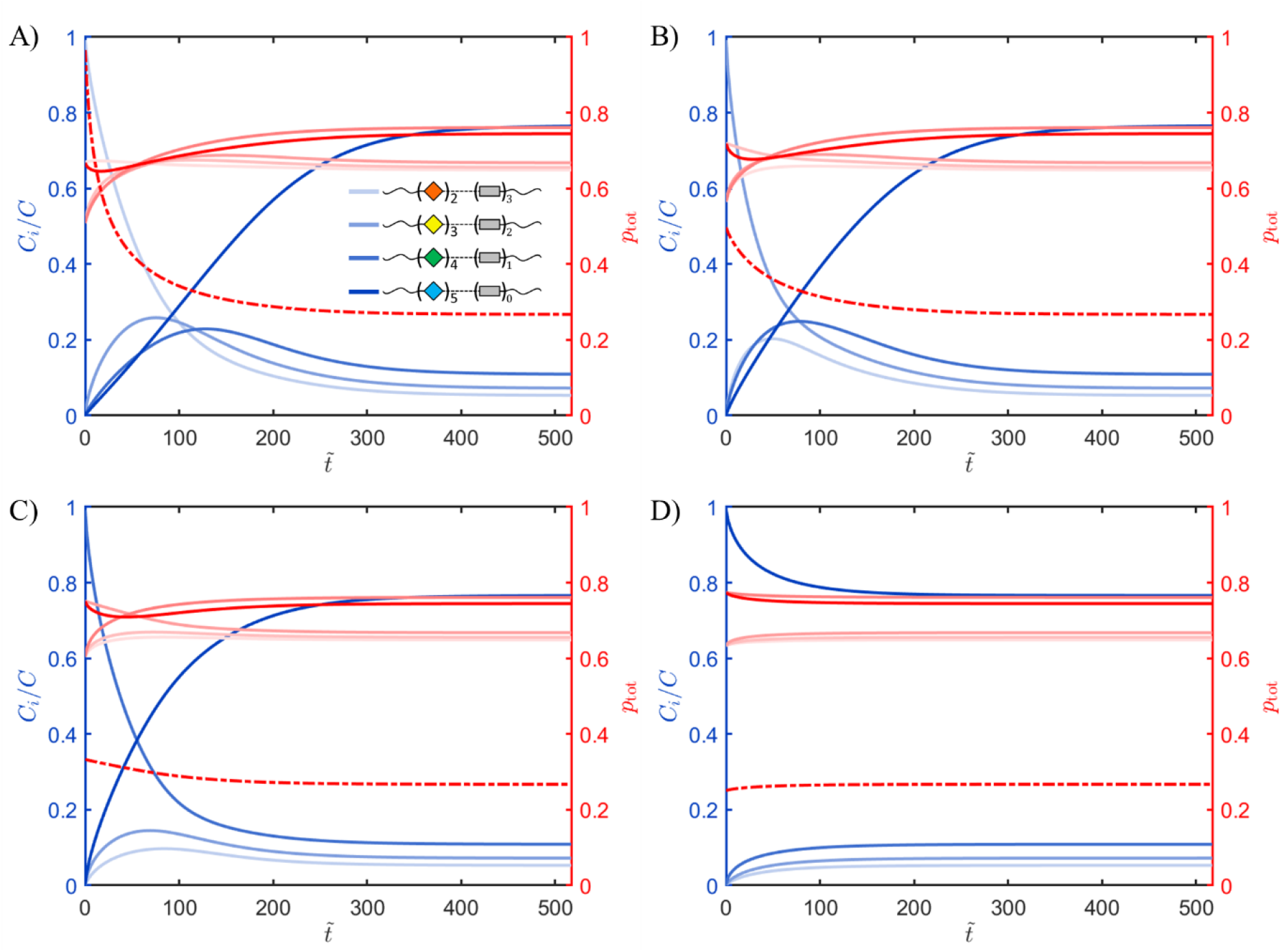
Valency distribution as a function of time. Panels A) – D) represent kinetic traces assuming an initial single species with an active sticker valency that increases from *ξ*_1_ = 2 (Panel A) to *ξ*_4_ = 5 (Panel D). All other input parameters are the same as for Figure 5A (see Section S6 in the SI). The plots show the concentration of IDP species *C*_*i*_ with different sticker valencies (see legend in Panel A), normalized by *C*_con_ (blue curves) and individual (red, shades) and total (red, dark) fractions of bound stickers, all plotted as a function of dimensionless time 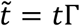. The dash-dotted red curves represent the mean-field percolation threshold.

In the next set of calculations (Figure 7) we show how the kinetics change by increasing the initial valency as *i.e*. 3 ≤ *ξ*_1_ ≤ 5, for similar number of species (*n* = 4), binding strength (*K* = 1024 M^−1^) and solvency (*χ*_*iS*_ = 0.68). Now we let the maximum valency scale with the number of species, so the most abundant component at 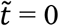 is the one with the lowest valency in the distribution at any time. have *p*_tot_(*t* = 0) > *p*_*g*_(*t* = 0), implying that a percolating network already exists at *t* = 0. These calculations reveal several other interesting properties. First, since the concentration of stickers increases with *ξ*_1_, the system tends to more strongly favor the highest valency species, with the transition between the transient and equilibrium regimes becoming more abrupt. Second, the time to reach equilibrium is longer if the valency is low, due to overall lower chemical potentials and hence lower driving force to generate new stickers through folding. Third, the minimum in the total bound fraction *p*_tot_ (dark red curve) shifts to later time and, finally, the occurrence of the peak in the concentration of species with intermediate valency, *i.e. ξ*_2_ and *ξ*_3_ (see blue arrows), coincide for higher total valency, probably because the fractional increase in the number of stickers per chain decreases and the species behave more similar.

**Figure 7.**
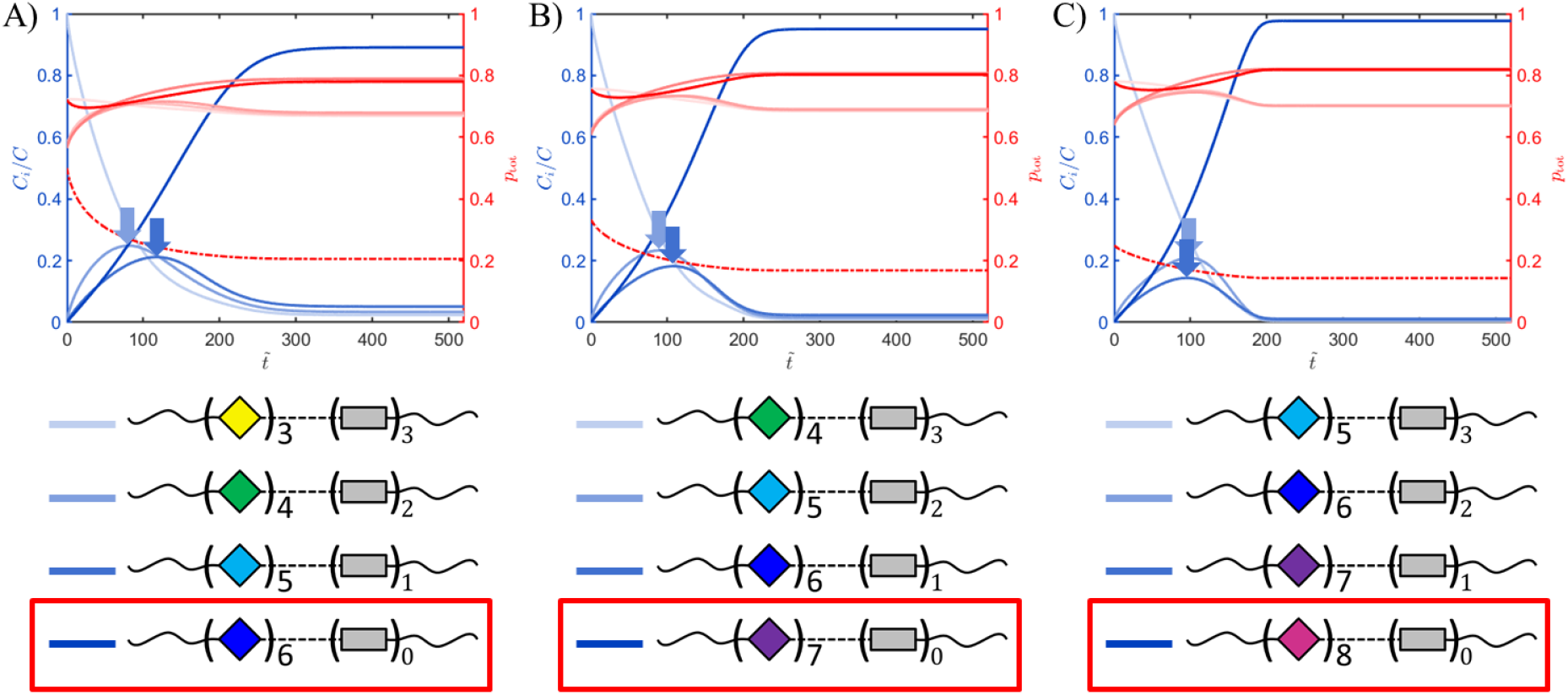
Aging kinetics in IDP-rich phase separated droplets as a function of minimum valency and constant number of species *n* = 4. Normalized concentrations are plotted as a function of dimensionless time for *χ*_*iS*_ = 0.68, *K* = 1024 M^−1^ and Δ*ω*_0_ = 0.0032 for the series of species given under each panel. The dash-dotted red curve represents the mean-field percolation threshold.

Since biological aging is subject to a ‘balancing act’ between relaxation towards equilibrium and prevention of that from happening,^[8]^ it is of interest to study how quickly an aging droplet or solution reaches (or tries to reach) equilibrium as a function of molecular properties. In the next set of calculations, we do this by tracking the aging kinetics as a function of the driving force by systematically changing the ‘conformational’ penalty Δ*ω*_0_ of sticker formation against the energy gain associated with sticker binding. We fix the minimum valency and the number of possible species at *ξ*_1_ = 2 and *n* = 7, *i.e*. representing an IDP that accommodates up to *ξ*_*n*_ = 8 stickers per chain. To illustrate the presence of a scaling, we assume a higher sticker binding constant than in the previous section. A sufficiently poor solvency guarantees that Scenario I applies: with *χ*_*iS*_ = 0.84 we obtain with the present parametrization (see Section S6 of the SI) a condensate concentration of *C*_con_ = 2.26 mM for *ξ* = *ξ*_1_ = 2, which increases negligibly by ~3% for *ξ* = *ξ*_*n*_ = 8.

Figure 8 shows the normalized concentrations, total bound sticker fraction *p*_tot_ and mean field percolation threshold *p*_*g*_ as a function of dimensionless time and for an increasing conformational energy penalty Δ*ω*_0_ (see caption). Again, we observe a number of interesting phenomena. Owing to the high value of the product *K* × *C*_con_ practically all stickers are in a bound state across the full-time domain, irrespective of the value for Δ*ω*_0_ (see solid red lines). If Δ*ω*_0_ is low, equilibrium is reached fast, with the highest valency species, *ξ*_7_ = *ξ*_*n*_ = 8, being strongly preferred. As Δ*ω*_0_ increases, the time to reach equilibrium increases and even diverges for a value 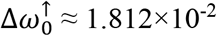, so somewhere between panels E and F. For 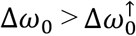 a second regime is observed, wherein the time to reach equilibrium seems to *decrease* with increasing Δ*ω*_0_ and where the species with the lowest valency, *ξ*_1_ = 2, is preferred.

**Figure 8.**
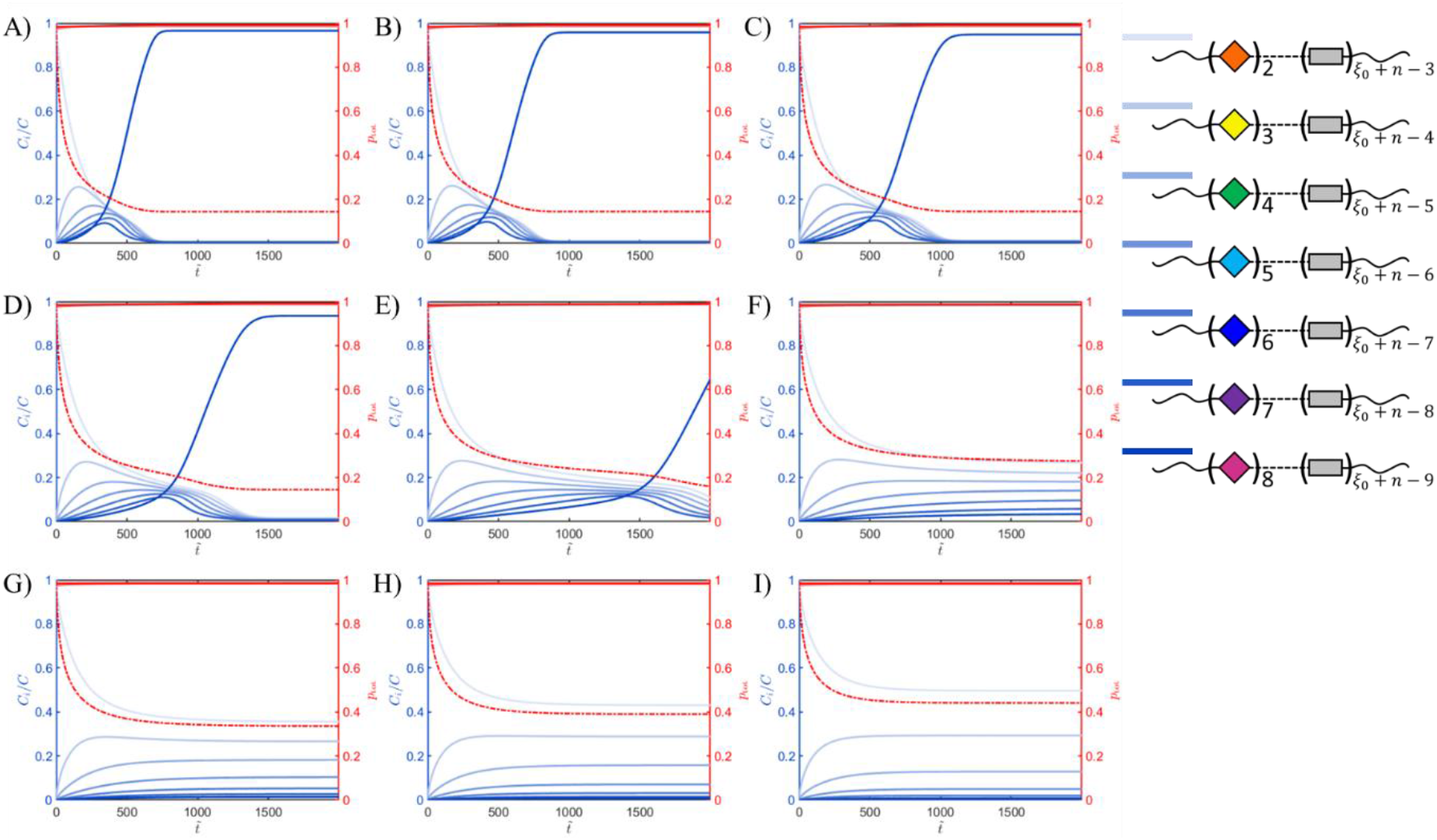
Aging kinetics as a function of the sticker formation energy. Normalized concentration of species with different valencies plotted as a function of dimensionless time 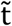 (blue curves, see legend) for *χ*_*iS*_ = 0.84, *K* = 528244 M^−1^, *ξ*_1_ = 2 and *n* = 7. The dimensionless conformational energy penalty Δ*ω*_0_ increases linearly from 1.72×10^−2^ in panel A to 1.88×10^−2^ in panel I in steps of 2.0×10^−4^. The solid and dash-dotted red curves respectively represent the total fraction of bound stickers 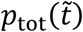 and the mean-field percolation threshold 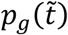.

The change in preference for the highest to the lowest valency species is explained as follows. When Δ*ω*_0_ increases, the cost of forming new stickers changes from being outweighed by the gain associated with sticker binding to outweighing the latter. Interestingly, the switch between these regimes is rather abrupt and ‘binary’: the system switches between *ξ*_1_ = 2 and *ξ*_7_ = 8 as the preferred valency at equilibrium. A range for Δ*ω*_0_ for which intermediate valencies are preferred is very narrow. Furthermore, if *K* × *C*_con_ is high and 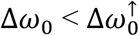, the shape of the trace for the highest valent species (here *i* = 7) becomes pronouncedly sigmoidal. If Δ*ω*_0_ is near, but just below 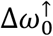, the concentration of *ξ*_7_ remains suppressed in a “pseudo-equilibrium” characterized by near zero net fluxes for a long time before suddenly rising towards its equilibrium which exceeds the concentrations all lower valency species. We note that the above implies that for a given concentration and binding strength, 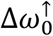 should increase with the maximum valency since sticker formation and association counteract each other energetically.

Irrespective of the value for Δ*ω*_0_, the chains form a percolating network (*p*_tot_ >> *p*_*g*_, solid and dash-dotted red curves) at early times, implying that in practice, already early on during aging IDP condensates are expected to be highly viscous and pronouncedly viscoelastic. The architecture of the network *does* however depend on the value of Δ*ω*_0_. To demonstrate this, we compare the equilibrium condensates obtained for a low (panels A – E) and a high (panels F – I) Δ*ω*_0_. For 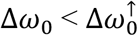 the network comprises virtually a single species (*ξ*_7_ = 8), of which all eight stickers are in a bound state. This implies that if the precursor sites are distributed evenly along the chains, the mesh is not only dense, but also regular in the sense that the strand length between crosslinks is constant everywhere. In contrast, if 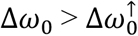, sticker valency is distributed, especially if Δ*ω*_0_ exceeds 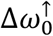 only moderately (panels F and G). As a consequence, even if the precursor sites are evenly distributed, the network will, on average, be more open and rather irregular due to significant variation in the strand length between crosslinks. In Section 2.5 we will showcase how the model can, in a crude way, estimate the actual viscoelasticity of the droplets using an approximate model based on a multi-component-multi-valency extension of ‘sticky Rouse theory’.

### 2.4. Experimental validation

Before calculating constitutive properties, we demonstrate that the above predictions in principle agree well with experimental *in-vitro* aging of fluorescently labelled FG nucleoporin-like IDP constructs based on the “perfect repeat” version of Nup98. As mentioned in the introduction, these comprise a 52-times repeat of the GLFG-based 12 amino acid sequence GGLFGGNTQ**X**AT (see SI, Section S6) and form ideal model systems to design and study IDP biophysics.^[33], [44]^ Besides a wild-type protein (GLFG-WT), for which **X** = P, we engineered two mutants comprising 8 and 18 evenly spaced sequences having **X** = V, which we dub GLFG-V_*x*_, with *x* = 8 or 18. Following previous experiments that show that the substitution of prolines from inter-FG spacer sequences can lead to amyloid-like interactions,^[33], [62], [63]^ we chose valine, a non-polar hydrophobic and aliphatic amino-acid, aiming to increase the aging propensity in our *in-vitro* droplet experiments.

We prepared condensates of the GLFG-repeat domain constructs through *in-vitro* phase separation^[41]^ and studied the fluorescence recovery dynamics by recording FRAP, at *t* = 1h, 24h, 48h and 72h after droplet formation (details in SI, Section S7). Figure 9 shows all recorded FRAP traces, each representing an average over multiple droplets within the same sample. For each aging time we show the traces for multiple (typically three) samples to demonstrate the degree of reproducibility. Clearly, the curves corresponding to GLFG-WT and GLFG-V_8_ do not show a clear trend with aging time, whereas the ones for GLFG-V_18_ are consistent with a gradual but non-linear decrease in dynamics beyond experimental error, though reaching a saturation at ~48h.

**Figure 9.**
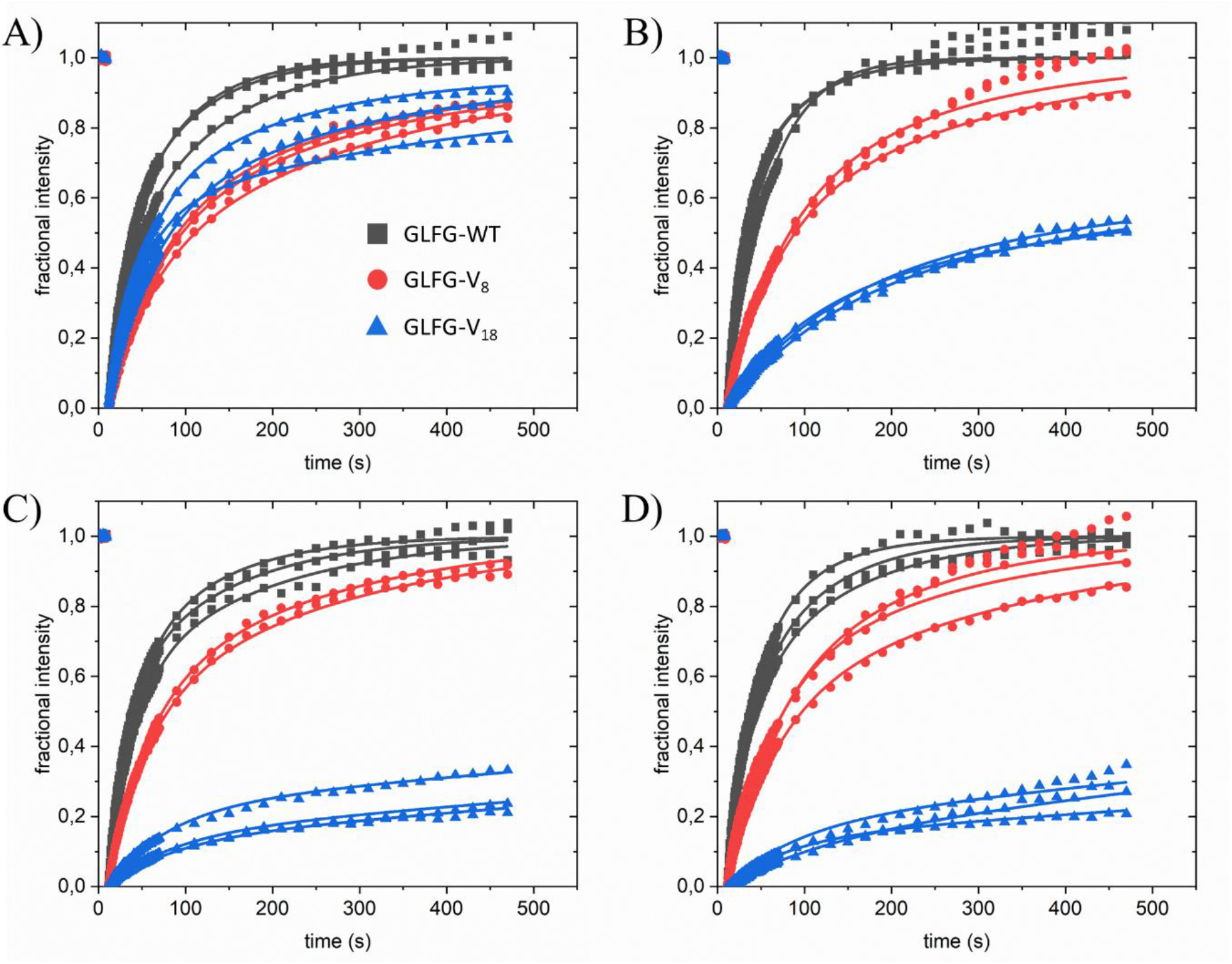
Measured fluorescence recovery traces (symbols) plotted as a function of aging times of 1h (A), 24h (B), 48h (C) and 72h (D). The x-axis in each panel represents the fluorescence recovery time within each FRAP experiment, whereas the aging time is encoded by the panel identity. The recovery traces were measured on phase separated droplets of the GLFG-repeat domain variants indicated in the legend in panel A. Traces with similar color within the same panel represent different samples to show experimental spread and reproducibility. The solid lines represent fits based on biexponential functions.

To get a general idea about (changes in) dynamics, we obtain for each trace in Figure 9 a fluorescence recovery half time (*t*_1⁄2_) by means of curve fitting on an empirical basis (see Section S7 of the SI for details). None of the curves fitted satisfactorily against a mono-exponential function, but all fitted quite well against a biexponential function. Some of the recovery curves, in particular for the GLFG-WT exhibited a drift beyond a full recovery of the fluorescence (see black curves in Figure 9B), which we ascribed to an experimental artifact. This was corroborated by the fact that omitting such drifts in the curve fitting gave consistent results in comparison with data that did not exhibit them. As explained in Section S7, the half times of separate experiments are averaged (Table S6), from which we calculate effective diffusivities as explained below.

The average recovery half times (see Table S6) are plotted as a function of aging time in Figure 10A. Clearly, only GLFG-V_18_ shows appreciable aging with dynamics saturating after about 48h (blue curve). In contrast, the wild type and V_8_ mutant do show some spread in the experimental value but no particular trend towards aging. However, GLFG-V_8_ (red) does consistently exhibit somewhat higher recovery times than GLFG-WT (grey). Overall, the experimental results reveal a strongly non-linear, perhaps critical dependence of the aging on the valency, as well as a generally higher stickiness of the GLFG-V_*x*_ mutants compared to the wild type.

**Figure 10.**
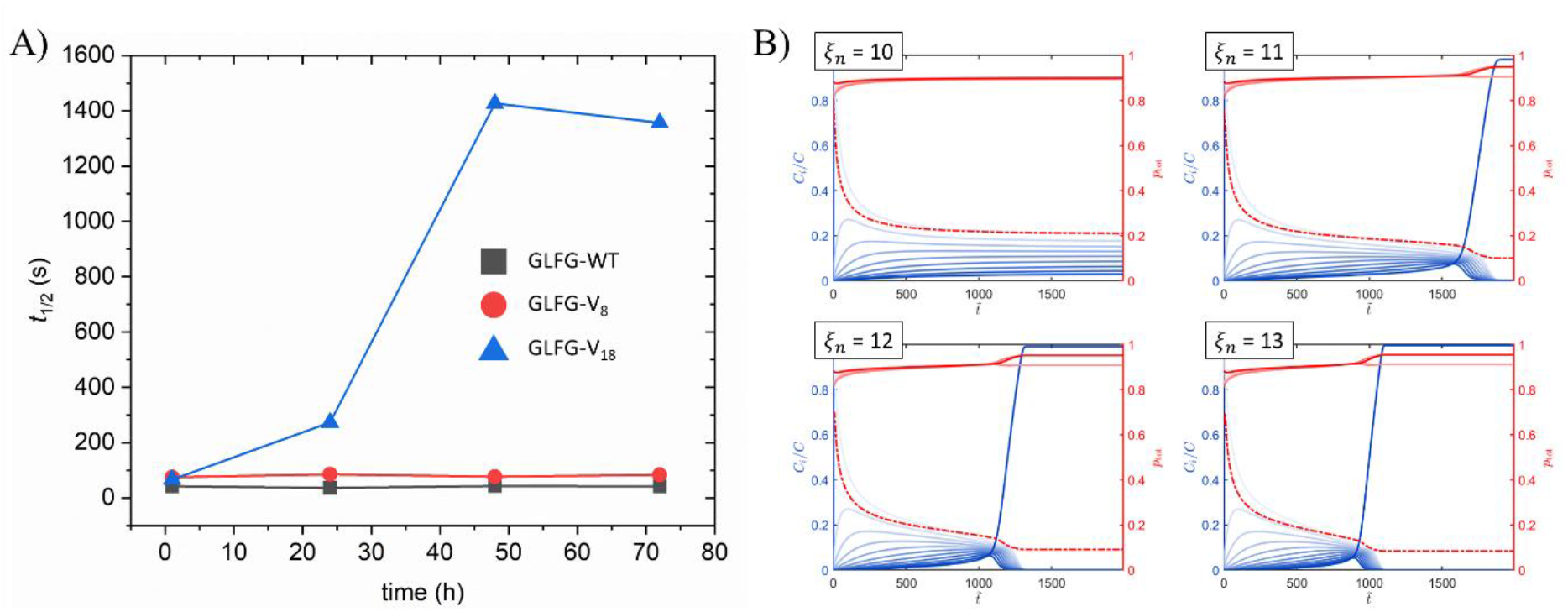
Comparison of experimental and computational results on aging as a function of time and valency. A) Mean fluorescence recovery half times (Table S3) measured with FRAP on labelled GLFG-repeat domain variants (see legend). B) Normalized concentrations (blue curves) of valencies plotted as a function of dimensionless time in phase separated droplets of an IDP with a maximum valency *ξ*_*n*_ as indicated above each panel and Δ*ω*_0_ = 2.50×10^−2^, *χ*_*iS*_ = 0.84 and *K* = 3072 M^−1^. Light to dark blue correspond to species with increasing valency. The solid red curves represent the fraction of stickers in a bound state on a per species (light) and total (dark) basis and the dash-dotted red curves represent the change in the percolation threshold with time.

Figure 10B shows the results of comparative model calculations. We studied a range of IDPs with increasing maximum valency *ξ*_*n*_ (as in the experiment) and calculate the (relative concentration of species with valencies 2 ≤ *ξ*_*i*_ ≤ *ξ*_*n*_ (blue curves) as a function of the dimensionless time passed after phase separation. The values for *ξ*_*n*_ are in principle chosen in the same range as the experimental valencies, although we are aware of the fact that it is uncertain how the number of P-to-V point mutations relates to an actual sticker valency. This does however not affect our conclusions in any way. Based on structural similarity, we assume the same solvency (*χ*_*iS*_ = 0.84) for all species and represent the 624 residues by *N* = 100 coarse-grained monomers, based on the notion that 6 – 8 residues have been identified to be the smallest sequence capable of undergoing an order-disorder transition.^[64]^ The sticker formation penalty and binding strength^[65]^ are fixed at Δ*ω*_0_ = 0.025, *K* = 3072 M^−1^. Starting with a single bivalent species (*ξ*_1_ = 2) at *t* = 0.

The calculations nicely reproduce the strongly non-linear dependence of the aging on valency observed for the GLFG-repeat domain variants: for *ξ*_*n*_ < 11 the valency distribution rapidly saturates with the lower valencies (lighter blue shades) remaining dominant at all times. Formulated as laid out in Section 2.3, this behavior is expected if the decrease in the total free energy owing to sticker association does not overcome the overall folding penalty. So, for low valencies we have 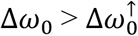. In contrast, for *ξ*_*n*_ ≥ 11 we have 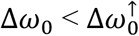 and sufficient complexes can form to sooner or later, depending on valency(!), overcome the sticker formation penalty. After a certain lag time, a steep rise is observed in the concentration of the species with the maximum valency (darkest blue shade in each panel), at which point the chains within the condensate are expected to become much less dynamic. This lag time decreases with *ξ*_*n*_.

In Figure 11 we compare this decrease in dynamics as suggested by the experiment and model. The figure plots the experimental (panel A) and computational (panel B) relative effective diffusivities 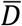 as a function of aging time, normalized by the effective diffusivity 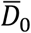 determined for the shortest interval, *i.e*. 1h in the experiments and 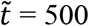 for the computations, so right after the steep initial drop in the concentration of the lowest valency species *n* = 1 with *ξ*_1_ = 2 (see Figure 10B). In a FRAP experiment, the effective diffusivity scales reciprocally with the fluorescence recovery half time as:^[66], [67], [68], [69]^ 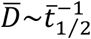, up to a multiplicative constant in units of m^−2^, which drops out by applying the same normalization as for the experimental data. This effective diffusivity obviously represents some sort of average over species with different valencies, as the exact composition of the mixture of chains in various folding states is not known.

**Figure 11.**
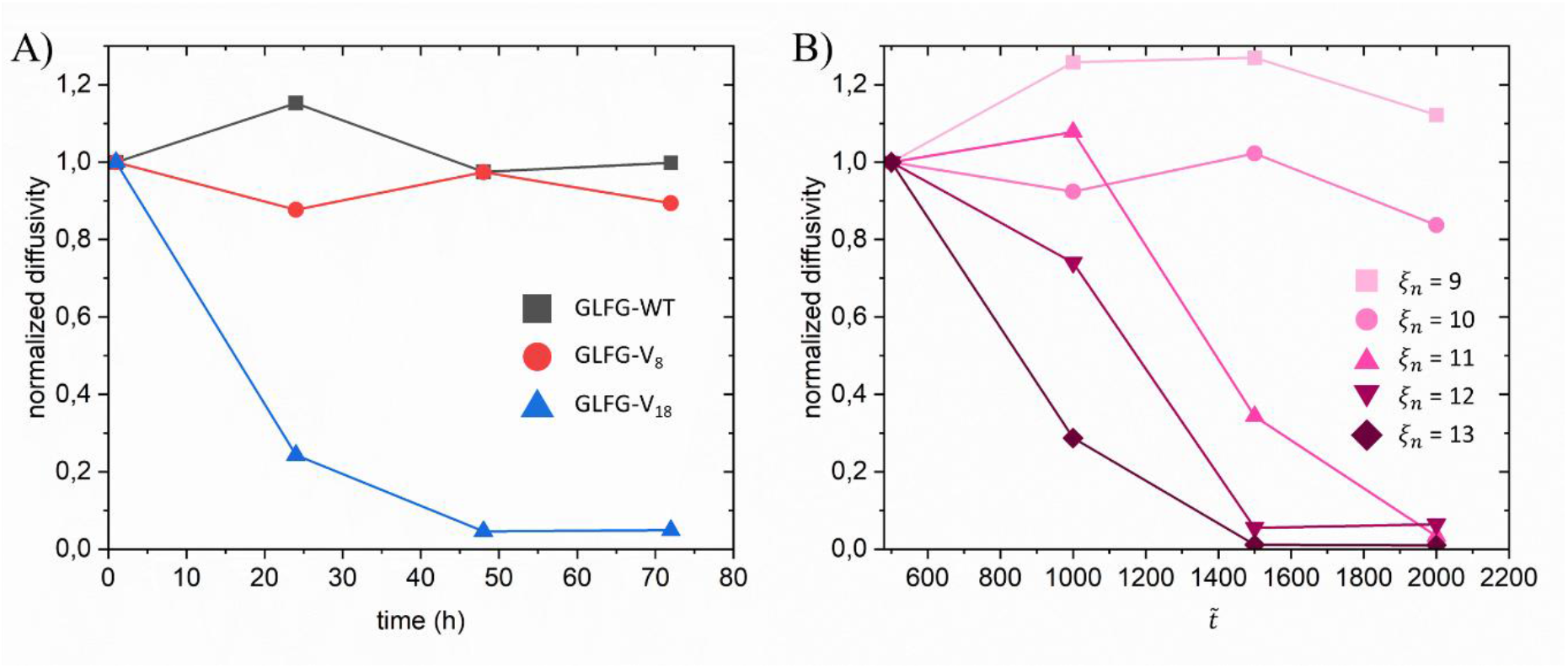
Experimental (A) versus modeled (B) normalized effective diffusivities in phase separated IDP droplets, plotted as a function of aging time. The latter is dimensionless in the modeled case. The legends show the GLFG-repeat domain variants (A) and maximum sticker valencies (B) of the curves in corresponding color.

In the same way, the computational effective diffusivities are averages of the values corresponding to the individual species within the valency distribution: 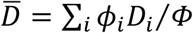. We calculate the latter using a multi-component version of the ‘sticky Rouse model’.^[70], [71], [72]^ Within this model, the monomeric friction coefficient *ζ*^(*i*)^ may be defined in two different ways, depending on whether the monomer is non-sticky or sticky. In case it is non-sticky, we use the Einstein-Smolukowski equation to define *ζ*^(*i*)^ = *ζ*_0_ = *k*_*B*_*T*⁄*D*_0_, with *D*_0_ the self-diffusivity of a solvent molecule or non-sticky monomer, which for all calculations we fix at 10^3^ μm^2^/s, *i.e*. roughly the self-diffusivity of water at room temperature.

In case the monomer is sticky, we have 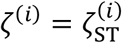, with 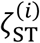 a rescaled friction coefficient of a sticky bead of IDP species *i* with valency *ξ*_*i*_. Since the number of stickers per chain is low compared to the number of non-sticky monomers, we follow Ricarte *et al*. by approximating the total friction a chain experiences by a linear combination of the contributions from the sticky and non-sticky monomers.^[70]^ We approximate 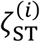 by a modified Einstein-Smoluchovski equation, which we derive in Section S3.1 of the SI:

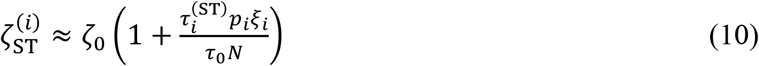

where *p*_*i*_*ξ*_*i*_ is the effective number of stickers in a bound state per chain of species *i* and 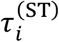 the residence time of a sticker of species *i* in a complex, which is related to 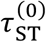 the bare life time of a sticker complex: 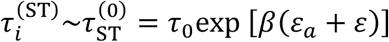 and further detailed. The Rouse diffusivity of species *i* is obtained as:

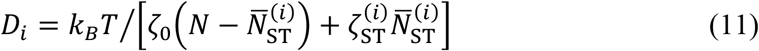

where the denominator represents the total friction coefficient of the chain, with 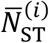 the expected number of non-covalent crosslinks per chain of species *i*.

The fact that *p*_*i*_ is concentration dependent and different for each valency implies that the friction due to stickers depends on dilution and valency. As explained below in Section 2.5, for dense networks, the residence of a sticker in a complex exceeds the bare life time due to repeated (re)formation of the same sticker complex In that case, the friction coefficient of a sticky unit *j* on species *i* becomes renormalized as: 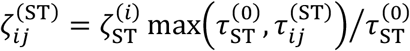, with 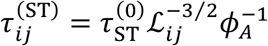 the effective residence time in the complex and ℒ_*ij*_ an effective interaction radius length and *ϕ*_*A*_ the volume fraction of free stickers. We refer to Section S4 of the SI for details. In case of renormalization, the total friction is obtained by summation of all individual coefficients.

In agreement with the experiments, the calculated results show that for low valencies the effective diffusivity does not depend on the aging time, whereas for higher valencies a stark decrease in dynamics is observed. To demonstrate the non-linearity more clearly, we included an additional IDP, with a maximum valency of *ξ*_*n*_ = 9 (see legend). The agreement between experiment and model is more subtle than just expressed by the overall trend. Namely, it is also encountered in the loss in the stochastic nature of the data if the valency becomes high. The randomness in the experimental data is likely introduced by a combination of sample inhomogeneity and handling, whereas in the computational case it is (deliberately) introduced by choosing the order of the conversion of the precursor sites into active stickers (see Section 2.5) randomly for each time point. With increasing valency, both the experiments and the calculations show that the spread that results from this randomness becomes more strongly subdued by the drop in dynamics, with a concomitantly clearer trend with aging time.

Our calculations assign a saturation of the dynamics with aging time (see curves for *ξ*_*n*_ = 12 and *ξ*_*n*_ = 13 in Figure 11B) per definition to an approach to an equilibrium, whether stable or metastable. This *may* be the case in the experiment as well, for instance if a highly viscous but still liquid non-amyloid aged state is reached that is metastable relative to a solid-like amyloid state. Sticker binding is reversible and the residence time in a sticker complex is not infinite, even for very high association constants. This means that in the limit of very slow deformation, the solution or condensate will still behave as a liquid. One could argue that this is also the case in any biological material we consider ‘solidified’ on account of the formation of non-covalent (and hence reversible) interactions. Of course, a very long residence time of stickers in the bound state would shift the cross-over frequency of the loss and storage modulus in a Maxwell plot (see Section 2.5) to very low values, meaning that the solution is technically a liquid but would, for all intends and purposes, behave as a solid. In Section S8, we show that, given some caveats, our model is to some extent amenable to describing a ‘true’ dynamic arrest, in the sense that all dynamic processes halt before the systems reaches a global equilibrium.

### 2.5. Viscoelasticity

In this section we showcase that our model can be used to give estimates of the constitutive properties (viscoelasticity) of the droplets and how these may change with time. Calculating these properties is attractive, since experimental dynamic and constitutive data of biomolecular condensates is becoming more widely available, providing the opportunity for gaining physical insight on how (time dependent) constitutive properties relate to *in-vitro* aging kinetics and phase behavior. Nevertheless, we note that due to the absence of dynamic arrest in our description, some care must be taken with comparisons with data on extensively aged, fully solidified, condensates. In what follows, we will compare calculated dynamics with measurements on native and moderately aged IDP condensates.

To analyze how the viscoelasticity of a condensate depends on binding strength, solvency, valency distribution and time, we calculate its storage (*G*′(*ω*)) and loss (*G*′′(*ω*)) moduli as a function of an oscillatory deformation frequency *ω*. The moduli quantify to what extent the condensate is capable of storing and dissipating the energy associated with a deformation. In other words: to what extent they behave as an (elastic) solid or a (viscous) liquid. It is intuitive that for the present systems the viscoelasticity will not only be determined by trivial parameters, such as chain length and concentration, but particularly by the lifetime of sticker complexes *τ*_ST_, which are co-determined by the association strength.

Just as its classical predecessor,^[73]^ the sticky Rouse model ignores hydrodynamic interactions, which are indeed expected to be screened at concentrations as high as *C*_con_. Our approach, which we have introduced in Section 2.4 and detail in Sections S3 and S4 of the SI, generalizes the treatment by Ricarte *et al*. in that the effective friction coefficient 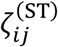 of a sticky monomer *j* of a species *i* is a function of *concentration and time*, due to its dependence on the bound fraction *p*_*i*_(*t*). Our model furthermore generalizes the analysis by Semenov and Rubinstein^[15]^ in that 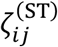 *varies among valencies* and, as explained below, may even differ *per sticker on the same chain*. The latter is related to the fact that, in contrast to the free energy, the sticky Rouse model is in fact sequence specific.

For sufficiently dense networks, Semenov and Rubinstein argued that the lifetime of a sticker complex should be renormalized to account for the fact that the same pair of stickers repetitively recombines before actually relaxing stress when one of the pair’s members binds to a new partner (see Figure 12). In that case, the effective friction coefficients associated with stickers on the same a chain should be considered *separately* since, depending on the monomer and precursor sequence, each sticker has a different probing volume wherein the encounter with a new binding partner may take place. This probing volume is determined by an effective strand length ℒ_*ij*_, which depends on the association state of neighboring stickers on the same chain, as explained in Section S4 of the SI. If the non-covalent network contains multiple species with different valencies (as in Section 2.3), the variation in ℒ_*ij*_ is inherently large, which means that the rheology of the network may become sequence dependent (see below).

**Figure 12.**
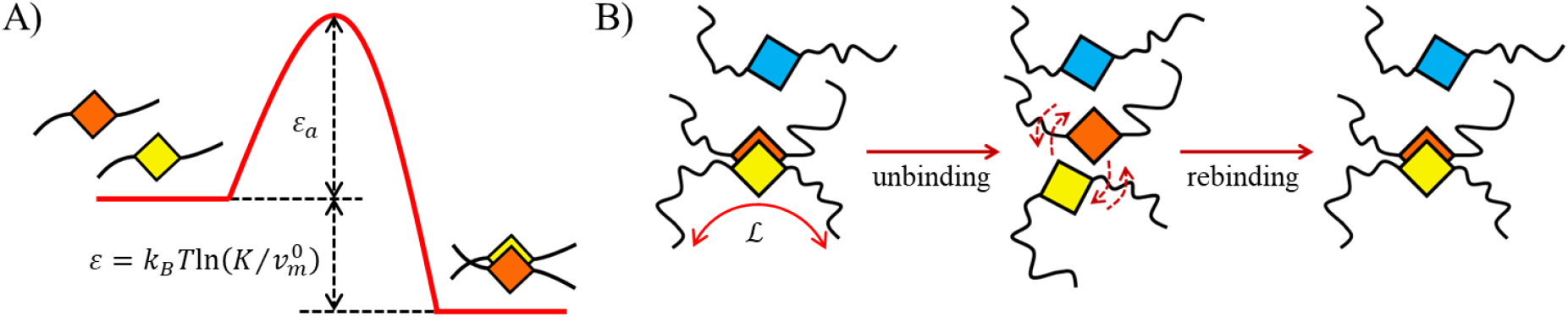
Schematic representation of the process of association and dissociation. A) Energy difference and barrier between the free and bound state. B) Unbinding and rebinding of the same sticker pair (orange and yellow) in the presence of a prospective new binding partner (blue). Inspired by Ref. [15].

We calculate the storage *G*′(*ω*) and loss 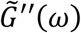 moduli using a generalized Maxwell model detailed in Section S3.2 of the SI and represented by Equations S18 and S19, taking into account the individual contributions of all species. Unlike more sophisticated treatments,^[74]^ our approach does not describe the dynamics associated with the breaking and formation of large clusters into/out of smaller ones^[15]^ and is therefore strictly valid sufficiently far above the percolation threshold, where the strand length between crosslinks is of order ~*N* or smaller. According to classical gelation theory, this is the case when (*p*_tot_⁄*p*_*g*_− 1)^−2^ < 1.^[15]^ Our kinetics traces show that owing to the high concentration inside the droplet, even for moderate binding strength *p*_tot_ is high and nearly time-invariant, whereas *p*_*g*_ initially drops very rapidly. This implicates that a strongly percolated regime is reached already early on in the aging process.

To study the influence of the topology of the network fluid, we calculate the effect of the sticker formation penalty on the viscoelasticity and the terminal viscosity of a phase separated droplet containing a heptavalent IDP with valencies as given in Figure 13. Within our model the order of precursor-to-sticker conversion does not affect the free energy and the aging kinetics, but is expected to influence the rheology of the condensate somewhat on account of the fact that the sticky Rouse model is in fact sequence specific. We calculate the viscoelastic moduli below and above the cross-over value 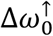, where the network topology supposedly changes drastically (see discussion above). The results of the calculations, as well as the other input parameters are given in the caption of Figure 14 and Section S6 of the SI.

**Figure 13.**
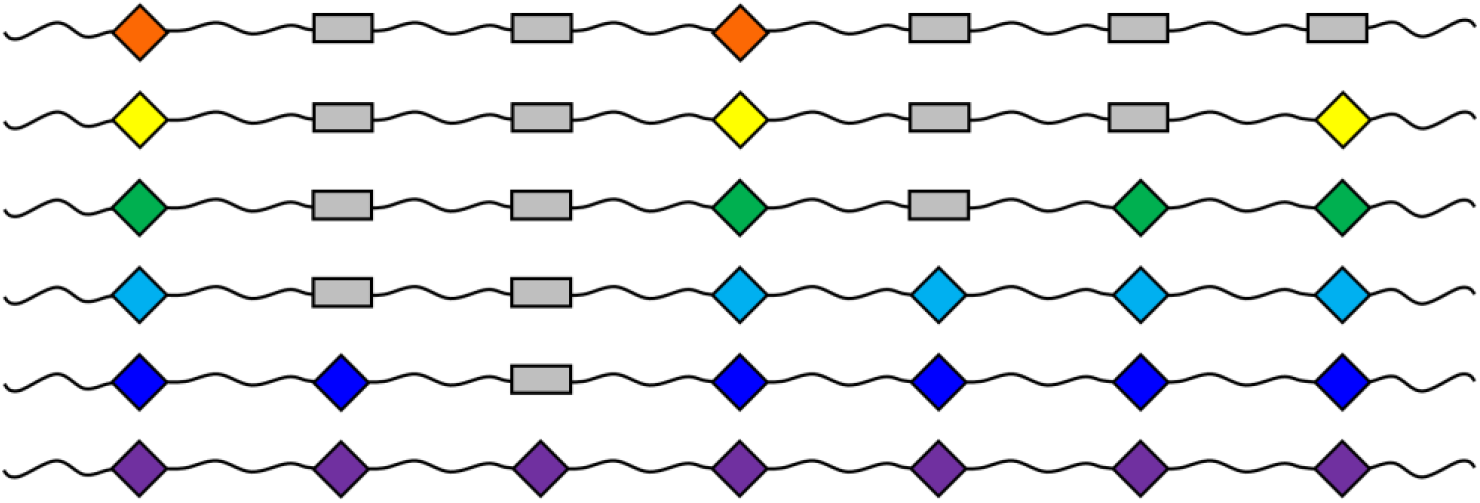
Schematic representation of the order in which precursors convert to stickers considering a heptavalent IDP in phase separated droplets of which we calculate the rheological properties below.

**Figure 14.**
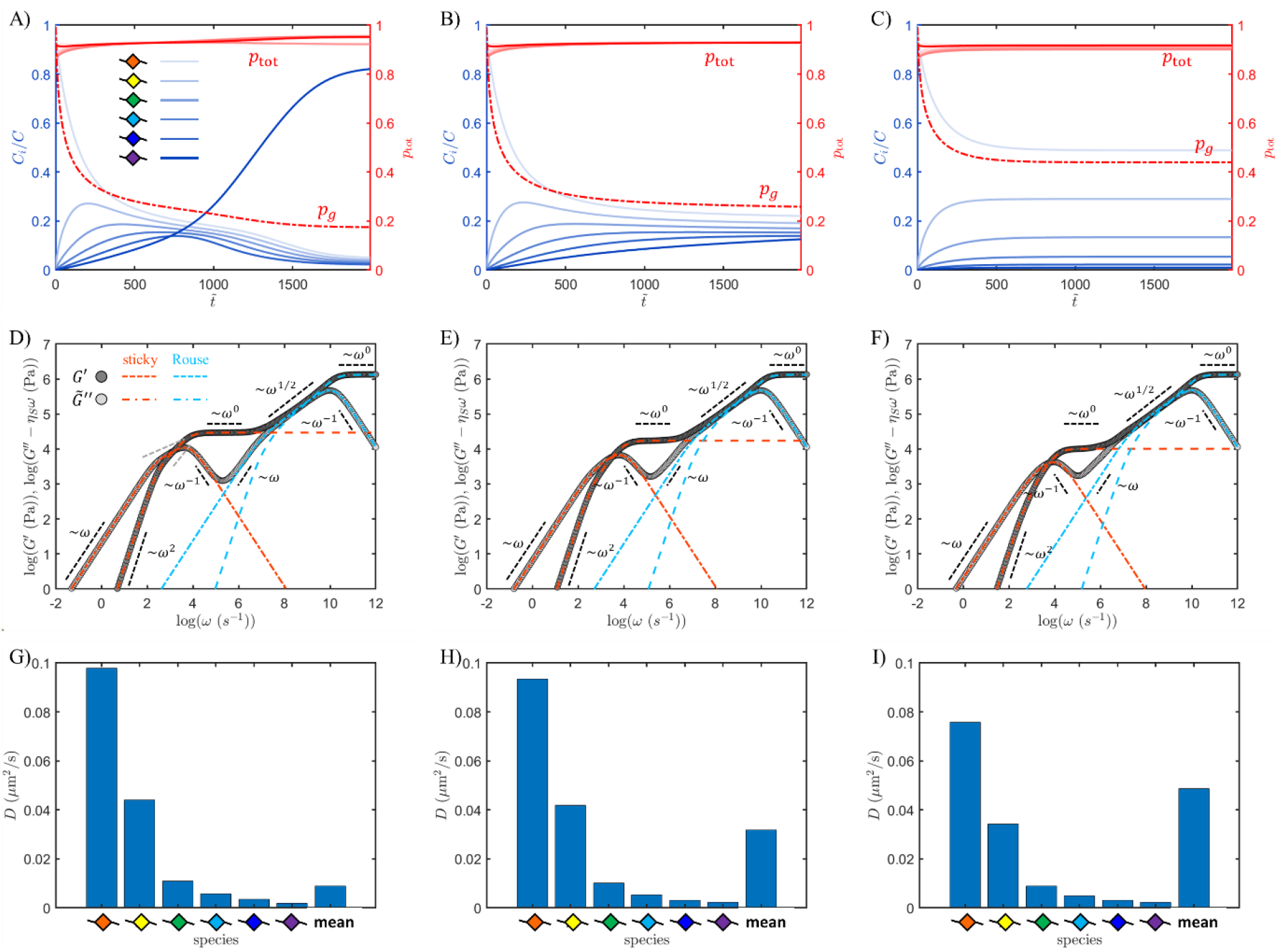
Aging kinetics, rheology and diffusivity of the heptavalent IDP depicted in Figure 13, calculated for a dimensionless sticker formation penalty of Δ*ω*_0_ = 1.10×10^−2^, 1.12×10^−2^ and 1.20×10^−2^ (left to right), with *χ*_*iS*_ = 0.84, *K* = 16384 M^−1^, *ξ*_1_ = 2 and *n* = 6. Panels A – C plot the normalized concentrations of the valencies as a function of dimensionless time (legend in A). Panels D – F plot the calculated storage (*G*′, dark grey symbols) and normalized loss modulus (*G*^′′^ − *η*_*S*_*ω*, light grey symbols) of the droplet at the final time as a function of frequency; the orange and blue dashed and dash-dotted lines represent the contributions to the moduli from sticky and Rouse models, respectively (see legend in D). Panels G – I plot the calculated individual and volume-averaged diffusivities 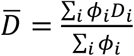, as indicated.

Figure 14 shows from top to bottom the kinetics traces, the viscoelasticity and the diffusivity of the species depicted above for, from left to right, increasing sticker formation penalty Δ*ω*_0_ (see caption) but constant association strength *K* and solvent compatibility *χ*_*iS*_ (see caption). The kinetics traces in Figures 14A – 14C exhibit a similar behavior as for the octavalent species discussed earlier: a low Δ*ω*_0_ leads to an eventual preference for the highest valency polymer, whereas a high penalty results in a dominance by the lowest valency species. As a result, with increasing Δ*ω*_0_ the network becomes less dense, expressed by a smaller difference between *p*_tot_ and *p*_*g*_ (solid and dashed red lines). Figures 14D – 14F shows the storage and loss moduli of the droplet in its final state as a function of frequency. We distinguish five dynamic regimes: with increasing deformation frequency we encounter i) a Newtonian regime 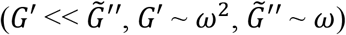, ii) a second liquid-like regime 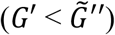 where the frequency scaling seems non-trivial and possibly non-universal, iii) a solid-like regime 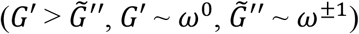, iv) a Rouse regime 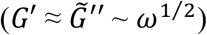 and v) a glassy regime 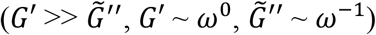. Encouragingly, the curves produced by our model are in general very good agreement with experimental ones, *e.g*. as obtained from unentangled melts of poly (ureidopyriimidione-*co*-acrylate) sticker-spacer polymers.^[75]^

The sticky- and Rouse-contributions to the viscoelastic moduli are given by weighted averages of the, respectively, *ξ*_*i*_ – 1 slowest and *N* – *ξ*_*i*_ fastest relaxation modes. Clearly, the first three regimes are dominated by sticker association. Under very slow deformation (regime i), the droplets are viscous liquids of which the (terminal) viscosity, given by 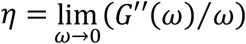, varies by an order of magnitude: *η* = 21, 6 and 2 Pa·s from low to high sticker formation penalty Δ*ω*_0_ (see SI, Section S5). These calculations reveal that high terminal viscosities are reached, even for a moderate binding strength and despite the assumption of purely binary sticker association rather than higher-order or fibril-like aggregation. For low Δ*ω*_0_ (panel D) up to the first cross-over between *G*′ and 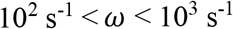 at 10^2^ s^−1^ < *ω* < 10^3^ s^−1^, a second, non-Newtonian, liquid regime (regime ii) appears, where the frequency scaling non-trivially depends on the network topology (dashed grey lines), with apparent slopes smaller than the typical values in the Newtonian regime. This regime becomes less prominent as the network becomes less dense for higher Δ*ω*_0_ (panels E an F).

At frequencies exceeding the first cross-over, deformation is fast compared to the complex life times. As a result, solid-like behavior is observed (regime iii), which, as expected, is more pronounced for a low Δ*ω*_0_, for which the network density is high: the cross-over frequency increases with Δ*ω*_0_, with the plateau in *G*′(*ω*) showing the opposite trend. Clearly, the droplets behave as viscoelastic fluids, which is in validatory agreement with experimental observations on a range of native or mildly aged biomolecular condensates.^[30], [43], [76], [77], [78]^ The present input gives values for the plateau modulus, cross-over frequency and terminal viscosity consistent with experimental data reported for condensates of the prion-like low complexity (PLC) domain of hnRNP-A1,^[43]^ as well as RG-rich low complexity IDP / dT40 RNA co-condensates.^[79]^ Naturally, for higher valencies and/or stronger association, more solid-like behavior and higher viscosities are expected, consistent with higher levels of maturation. Frequencies exceeding ~10^7^ s^−1^ (regime iv) address non-associated chain segments that do not “know” that they are part of a network, giving rise to the typical Rouse scaling until the deformation rate exceeds the relaxation rate 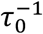 of a non-sticky monomer, marking the transition to a glassy regime (regime v). Also this is in agreement with experiments: The Rouse scaling following the viscoelastic plateau has for instance been observed in the rheology of melts of associating polymers.^[75]^

Figures 14G – 14I show the tracer diffusivity of each species *i* within the phase separated droplet, obtained as: 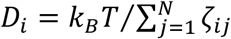, with *j* counting monomers and *ζ*_*ij*_ the friction coefficient of the *j*^th^ monomer of species *i* (see Section 2.4). For the given input (see caption Figure 14), we find diffusivities in the range 10^−3^ – 10^−1^ μm^2^/s, depending on valency and degree of association. These numbers compare favorably with those typically encountered for IDPs inside condensates and are consistent with an enhanced ‘stickiness’ consequential to an IDP’s capability of forming strong intermolecular bonds upon conformational adaptation (as in our calculations): diffusivities of the order ~10^−2^ μm^2^/s have for instance been measured for IDPs recruited into condensates of tumor suppressor protein p53, exhibiting tight interactions with the host through conformational changes.^[80]^ These diffusivities are about a factor ten lower than measured for folded proteins within the same host and that lack the capability of adapting their conformation.

The calculated diffusivities are furthermore consistent with those measured for FUS-LCD in condensates (~10^−1^ μm^2^/s),^[67], [81]^ also believed to be dominated by enhanced intermolecular interactions. Note with this respect that the diffusivities of non-phase separated IDPs in the crowded (! cytoplasm, where such ‘induced-fit’ interactions are absent, are typically three to even four orders of magnitude higher.^[82]^ The favorable comparison between the experimental works and the above calculations demonstrates that the magnitude of the strength of specific interactions responsible for the stickiness of conformationally adaptable IDPs inside liquid condensates relies on specific interactions with moderate association strength of the order ~10^4^ M^−1^. Finally, we note that the fact that the tracer diffusivity of the low valency species (*e.g. ξ* = 2) is highest for the highest network density (*i.e*. lower Δ*ω*_0_) is a consequence of its very low concentration (Figure 14A and 14G), due to which its stickers are mostly non-associated.

Below we demonstrate the model’s capability, although crude, to predict the development of the viscoelasticity in time. For this, we calculate the moduli at three time points during the aging process for the droplet corresponding to Figure 14A (low Δ*ω*_0_). For each of these points, marked by the dark orange arrows in Figure 15A, we ensure that (*p*_tot_⁄*p*_*g*_ − 1)^−2^ < 1 to principally obey the dense-network criterium. Figure 15B overlays the moduli calculated for each time point, again with delineated sticky and Rouse contributions. One can clearly discern the increase in plateau modulus (stiffening of the gel-like state) as well as the decrease of the first cross-over frequency, indicated by the black arrow. As shown by the dash and dash-dotted blue curves, the model correctly reproduces the expectation that the Rouse modes do not change in time, since they are determined by the relaxation of spacer segments.

**Figure 15.**
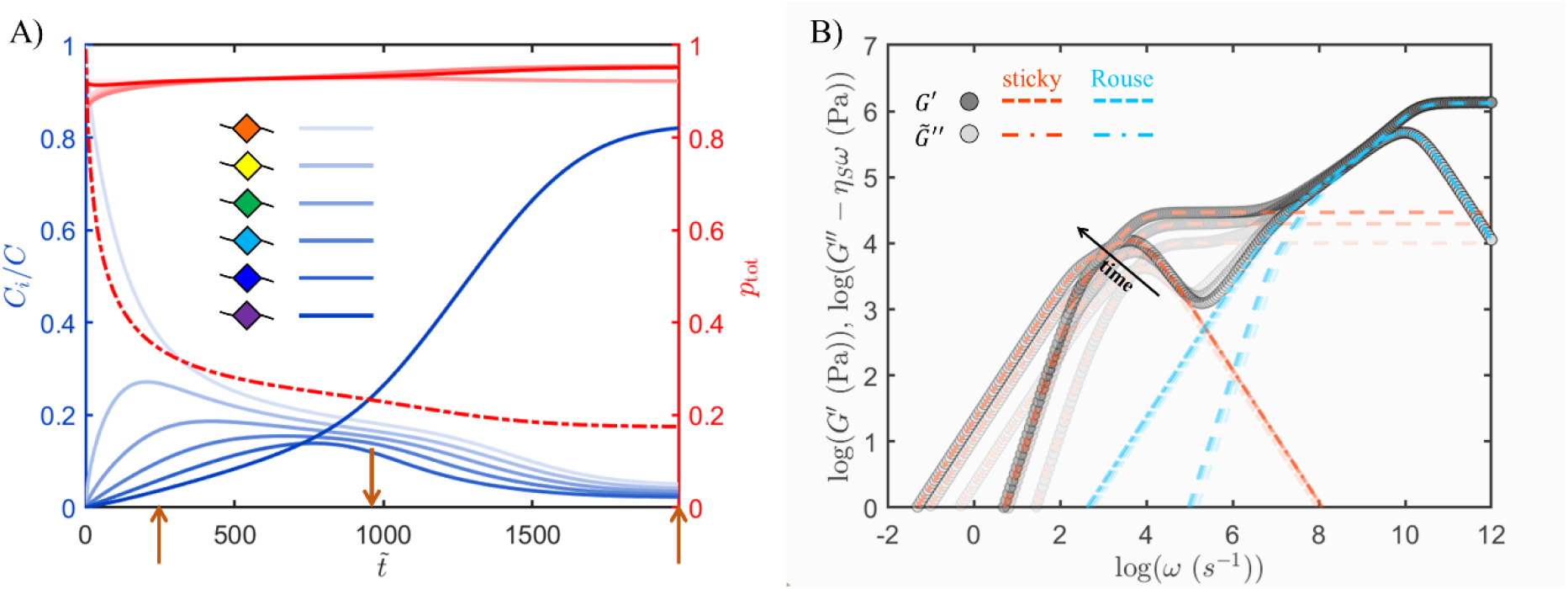
Aging kinetics and development of the rheology in time of the heptavalent IDP depicted in Figure 13, calculated for Δ*ω*_0_ = 1.10×10^−2^, with *χ*_*iS*_ = 0.84, *K* = 16384 M^−1^, *ξ*_1_ = 2 and *n* = 6 (so *ξ*_*n*_ = 7). Panel A (same as Figure 14A) plots the normalized concentrations of the valencies as a function of dimensionless time (see legend in A). Panel B plots the calculated storage (*G*′, dark grey symbols) and normalized loss modulus (*G*^′′^ − *η*_*S*_*ω*, light grey symbols) of the droplet at the aging times indicated by the dark orange arrows on the time axis in panel A as a function of frequency; the orange and blue dashed and dash-dotted lines represent the contributions to the moduli fromsticky and Rouse modes respectively (see legend in B).

## 3. Summary and Conclusion

Phase separation and condensate formation is omnipresent in biology and life science. Commonly, condensates become more viscous over time and can harden out in a process referred to as “molecular aging”. Our understanding of phase separation within the context of heteropolmers as IDPs in biology has grown dramatically over the past decade. However, kinetic models that intuitively capture the fact that most biological condensates become more viscous and solidify over time are scarce. This work addresses this with a time-dependent, multi-component extension of associating polymer theory that describes phase separation and aging of an IDP capable of recruiting binding partners through local, reversible folding within a unifying thermodynamic framework. The model demonstrates how the two phenomena are coupled and that the Second Law of Thermodynamics applies throughout, whether phase separation precedes and encourages aging or, *vice versa*, whether the increase in “stickiness” during aging drives phase separation. In our model “stickers” represent amino acid *sequences* that form through a reversible conformational change between a disordered and an ordered state, of which the latter may act as a binding partner for another such sticker. Hence, a “dormant” sticker can become “active” during aging, giving rise to a multicomponent solution of IDP chains with different valencies.

In the absence of an energy source, the chemical potential governs the transition between the dormant and active state, allowing us to study how both *in-vitro* phase behavior and aging are affected by solvency, concentration, molecular size, binding strength, valency, conformational energy penalty, etc. We achieve this by combining an extended free energy function with a set of coupled flux equations for the formation of each species. We identify two experimentally realistic scenarios for the phase behavior of the aging IDP: one in which phase separation largely precedes aging, but wherein aging is enhanced within the condensates and one wherein aging commences in the homogeneous solution and ultimately drives phase separation as well.

We show that the discriminator between these scenarios is the position of the initial (un-aged) state in phase space. The first scenario occurs if the initial state is already inside the miscibility gap, *e.g*. due to a poor solvency. Phase separation occurs fast before significant aging sets in. Aging then occurs exclusively within the dense phase. The second scenario is difficult to capture by means of a phase diagram. The initial position in phase space is typically near a critical point and, loosely formulated, finds itself inside a widening miscibility gap as more sticky interactions form. The model predicts aging to occur until an equilibrium is reached and is therefore particularly compatible with reversible β-sheet aggregation, but reproduces features characteristic to amyloid aging and is, preliminarily, amenable to include dynamic arrest. It predicts the state of folding in coexisting dense and dilute phases, as well as whether the bound sticker fractions either or not exceeds the percolation threshold. We show for the first scenario how the distribution of valencies of active stickers depends on the energy penalty for conformational change and identify a critical penalty above which the formation of high valency species is suppressed.

We validate the predicted steep valency-dependence of the aging kinetics using *in-vitro* phase separation and aging essays on disordered GLFG-Nup domain-like IDPs. We compare the 52 repeats sequence of the wild-type (WT) protein (proline present in the 52 spacers) with variants, wherein either 8 or 18 equally spaced proline residues from the inter-spacer region have been exchanged by a valine (V) residue. Using FRAP we studied changes in the dynamics of the IDPs inside condensates as a function of time. As predicted, we observe a very steep dependence of the aging kinetics on the sticker valency: whereas droplets of the GLFG-WT and the GLFG-V_8_ variant show no appreciable aging within 72h, GLFG-V_18_ droplets show a substantial drop in dynamics and achieving a highly viscous state after 48h. The semiquantitative agreement between experiment and model demonstrates that binary association with moderate strength can explain significant aging without having to consider the formation of an extensive fibrous amyloid network, which often develops on longer time scales.

In the final section, we demonstrate how the model can be used to estimate the viscoelasticity of aging condensates, while comparing our calculations with experimental literature. We apply the “sticky Rouse model” to calculate the storage and loss moduli as a function of deformation frequency. The model shows how the viscoelasticity of the condensate, as well as the diffusivity of the separate species, depend on the binding strength and network topology. We show that physically reasonable input for binding strength, valency and polymer size yields viscoelastic moduli and diffusivities that compare favorably with experimental data on various IDP condensates, as well as associating polymer melts. Lastly, we note that although our model predicts high viscosities and pronounced viscoelastic behavior, it does not capture ‘solidification’ as such, since binding sticker is always treated as reversible. Furthermore, in this work we have not addressed how the valencies appearing during aging distribute in space and thereby determine condensate heterogeneity. In future work we will address this by integrating the free energy functional and reaction flux equations with a full set of equations of motion for the polymer solutes.

## Acknowledgements

This project was funded by SFB 1551 “Polymer Concepts on Cellular Function” Project No. 464588647 and SPP 2191 Project No. 402723784 of the DFG (Deutsche Forschungsgemeinschaft). The authors acknowledge the IMB Microscopy Core Facility, supported by DFG funding under Project No. 497669232.

## Data Availability Statement

((include as appropriate, including link to repository))

Received: ((will be filled in by the editorial staff))

Revised: ((will be filled in by the editorial staff))

Published online: ((will be filled in by the editorial staff))

## Supporting Information

Supporting Information is available from the Wiley Online Library or from the author.

**Section S1**. Sticker free energy

**Section S2**. Binding equilibria

**Section S3**. A multicomponent sticky Rouse model

**Section S4**. Effective strand length and sticker life time renormalization

**Section S5**. Terminal viscosity

**Section S6**. Input parameters

**Section S7**. Phase separation and aging essays of perfect repeat nucleoporin constructs

**Section S8**. Dynamic arrest

