## Supplementary Information for "A Unifying Thermodynamic Model for Phase Separation and Aging of Biopolymers"

**Section S1. Sticker free energy**

By extension of the original associating polymer model for a homosolution,[1] We write the partition function for a solution containing $n$ associating polymers in the absence of nonspecific exchange interactions as:

$$Z = Z_{\text{ideal}} P_{\text{comb}} W \exp\left(-\frac{1}{2}\sum_{i=1}^{n}\sum_{j=1}^{n}\left(\left(1+\delta_{ij}\right)\frac{m_{ij}u_{ij}}{k_B T}\right)\right) \tag{S1}$$

Here, $Z_{\text{ideal}}$ is the ideal gas partition function and $m_{ij}$ is the number of a binary $ij$-sticker complex with binding energy $u_{ij}$. The factor $1/2$ in front of the double sum in the argument of the exponential prevents double counting of the number of heterotypic complexes and the Kronecker delta $\delta_{ij}$ prevents under-counting the number of homotypic complexes. The multicomponent combinatorial factor $P_{\text{comb}}$ becomes:

$$P_{\text{comb}} = \prod_{i=1}^{n}\prod_{j=1}^{n}\left[\frac{1}{2^{\frac{1}{2}(1+\delta_{ij})m_{ij}}\left(m_{ij}^{\frac{1}{2}(1-\delta_{ij})}\right)!}\right]\prod_{k=1}^{n}\left[\frac{m_{\text{ST}}^{(k)}!}{\left(m_{\text{ST}}^{(k)}-2m_{kk}-\frac{1}{2}\sum_{l=1}^{n}(1-\delta_{kl})m_{kl}\right)!\,m_{kk}!}\right] \tag{S2}$$

, with $m_{\text{ST}}^{(k)}$ total number of $k$-stickers. $W$ is the probability that all stickers involved in binding can be found close enough to their partners to form bonds in the absence of attractive forces:

$$W = \prod_{i=1}^{n}\prod_{j=1}^{n}\left(\frac{v_b^{(ij)}}{V}\right)^{\frac{1}{2}(1+\delta_{ij})m_{ij}} \tag{S3}$$

, with $V$ the total volume and $v_b^{(ij)} = x_{ij}v_0$ the volume of an $ij$-bond and $v_0$ the volume of a Flory-Huggins lattice site. Proceeding the usual way by taking the natural logarithm of the partition function and introducing Stirling's approximation yields an expression for the sticker contribution to the free energy density of the solution:

$$\frac{f_{\text{ST}}}{k_B T} = -\frac{1}{V}\ln\left(\frac{Z}{Z_{\text{ideal}}}\right) = -\sum_{i=1}^{n}\left[\frac{p_{ii}\phi_i}{2\bar{L}_i}\ln\left(\frac{x\phi_i}{e\bar{L}_i}\right)+\sum_{j=i+1}^{n}\frac{p_{ij}\phi_i}{\bar{L}_i}\ln\left(\frac{x\phi_j}{2e\bar{L}_j}\right)\right]+\sum_{i=1}^{n}\left\{\frac{\phi_i}{2\bar{L}_i}\left[p_{ii}\ln p_{ii} + 2\sum_{j=i+1}^{n}\left(p_{ij}\ln p_{ij}\right)+2\left(1-\sum_{k=1}^{n}p_{ik}\right)\ln\left(1-\sum_{k=1}^{n}p_{ik}\right)\right]\right\}-\frac{1}{2}\sum_{i=1}^{n}\sum_{j\geq i}^{n}\frac{u_{ij}p_{ij}\phi_i}{(1+\delta_{ij})\bar{L}_i} \tag{S4}$$

Here, $p_{ij}$ is the probability that an $i$-sticker is involved in a complex with a $j$-sticker. The value of $x = x_{ij}$, *i.e.* assumed the same for each sticker complex, becomes absorbed by the association constant according since: $K_{ij} = \frac{1}{2}xv_m\exp\left(u_{ij}\right)$, with $v_m$ the molar volume of lattice sites. Applying the binding equilibrium condition $\frac{\partial f_{\text{ST}}}{\partial p_{ij}} = 0$ reduces Equation S4 to the fourth term on the RHS of main text Equation 1.

**Table S1.** Input parameters for the calculation of the sticker free energy plotted in Figure S1.

| | | | |
|---|---|---|---|
| $N_1$ | 500 | $\phi_1$ | 0.12 |
| $N_2$ | 100 | $\phi_2$ | 0.30 |
| $N_3$ | 250 | $\phi_3$ | 0.18 |
| $\bar{L}_1$ | 20 | $\varepsilon_{11}/k_BT$ | 3.0 |
| $\bar{L}_2$ | 10 | $\varepsilon_{22}/k_BT$ | 5.0 |
| $\bar{L}_3$ | 15 | $\varepsilon_{33}/k_BT$ | 2.0 |
| $v_m$ (mol $l^{-1}$) | 1 | $\varepsilon_{12}/k_BT$ | 4.0 |
| $x$ | 2 | $\varepsilon_{13}/k_BT$ | 6.0 |
| | | $\varepsilon_{23}/k_BT$ | 3.5 |

To demonstrate that the model is internally consistent, we compare the sticker free energy of a ternary solution before and after enforcing binding equilibrium. More specifically, we calculate the dimensionless free density using Equation S4 and main text Equation 1 and plot the results in the same graph as a function of the binding probabilities $p_{ij}$ (see Figure S1). For these calculations we use an arbitrary set of input parameters for overall concentration, binding strength and valency as listed in Table S1. The equilibrium concentrations of complexes and unbound stickers, as well as corresponding probabilities $p_{ij}$, have been obtained according to the method given in Section S2. Substitution of the latter values in the fourth term on the RHS of main text Equation 1 yields the symbols in Figure S1. Independently varying each probability in Equation S4 while leaving the others constant yield the curves. The internal consistency of the model is demonstrated by the fact that i) in each case the equilibrium result coincides with the minimum in the free energy curve and ii) all minima correspond to the same free energy.

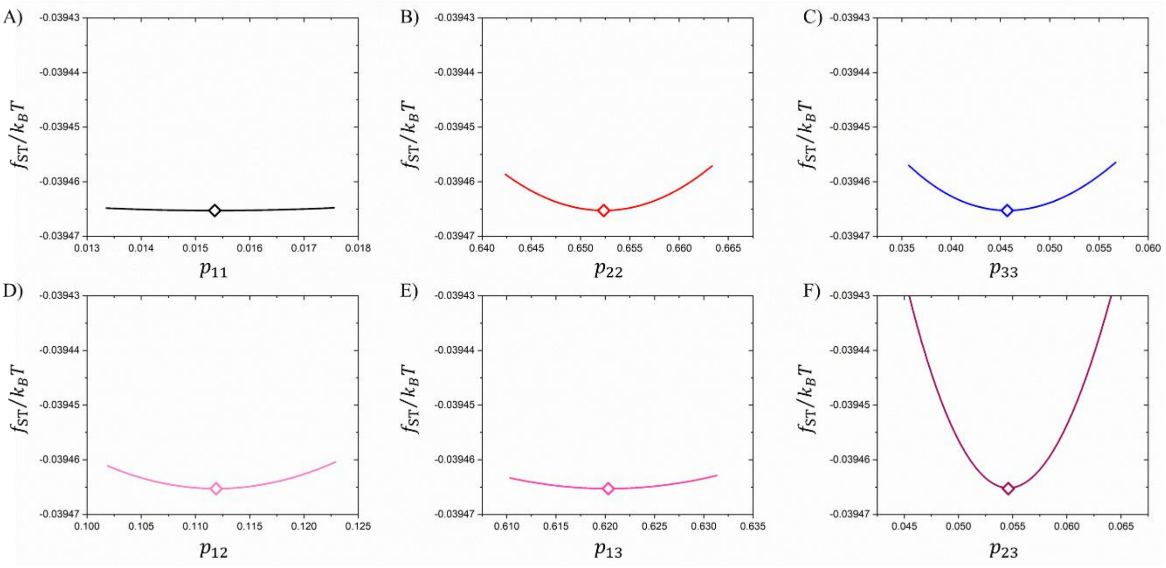


**Figure S1.** Dimensionless free energy density plotted as a function of the homotypic (top) and heterotypic (bottom) binding probabilities for a ternary solution. The lines and symbols have respectively been obtained using Equation S4 and the fourth term on the RHS of main text Equation 1 using the input parameters listed in Table S1.

**Section S2. Binding equilibria**

We consider a solution containing $n$ polymer species, each having a different valency. For each species, labeled $1 \leq i \leq n$ and molar concentration $C_i$ we solve the mass balance equation for the unbound sticker concentrations $[A_i]$, which gives the following set of target equations for a multivariate Newton-Raphson (NR) procedure:

$$\mathbf{g} = \begin{pmatrix} g_1 \\ \vdots \\ g_n \end{pmatrix} = \begin{pmatrix} [A_1]\left[1 + \sum_{j=1}^{n}\left(\left(1+\delta_{1j}\right)K_{1j}[A_j]\right)\right] - [S_1] \\ \vdots \\ [A_n]\left[1 + \sum_{j=1}^{n}\left(\left(1+\delta_{nj}\right)K_{nj}[A_j]\right)\right] - [S_n] \end{pmatrix} \tag{S5}$$

, with $[S_i] = [A_i] + \sum_{j=1}^{n}[B_{ij}]$ and $[B_{ij}] = K_{ij}[A_i][A_j]$ the molar concentration of binary sticker complex $B_{ij}$. The corresponding Jacobian matrix becomes:

$$\mathbf{J} = \begin{pmatrix} 1 + \sum_{k=1}^{n}(Y_{1k}[A_k]) & K_{12}[A_1] & \cdots & K_{1n}[A_1] \\ K_{21}[A_2] & 1 + \sum_{k=1}^{n}(Y_{2k}[A_k]) & \square & \vdots \\ \vdots & \square & \ddots & K_{(n-1)n}[A_{(n-1)}] \\ K_{n1}[A_n] & \cdots & K_{n(n-1)}[A_n] & 1 + \sum_{k=1}^{n}(Y_{nk}[A_k]) \end{pmatrix} \tag{S6}$$

, with $Y_{ij} = \left(1 + 3\delta_{ij}\right)K_{ij}$ an auxiliary variable and $\delta_{ij}$ the Kronecker delta. The vector of NR increments is obtained in the usual way according to:

$$\mathbf{d}[\boldsymbol{A}] = \begin{pmatrix} \mathrm{d}[A_1] \\ \vdots \\ \mathrm{d}[A_n] \end{pmatrix} = \mathbf{J}^{-1} \cdot \mathbf{g} \tag{S7}$$

, iteratively updating $[\boldsymbol{A}]_{t+1} = [\boldsymbol{A}]_t - \mathbf{d}[\boldsymbol{A}]$ and minimizing $||\mathbf{d}[\boldsymbol{A}]||$. Once all unbound sticker concentrations are known, the complex concentrations and bound fractions are trivially obtained, followed by three energies, exchange chemical potentials, reaction fluxes $j_i$ (see main text, Equations 3-5) and subsequently the polymer concentrations at the next time step. After that the above given NR procedure is repeated to equilibrate the sticker concentrations.

**Section S3. A multicomponent sticky Rouse model**

**3.1 Diffusion**

If the number of sticky monomers in a chain significantly exceeds the number of sticky monomers, the ratio of the diffusivity $D_R$ of a non-sticky and that of a sticky chain $D_i$ of species $i$ may in the Rouse limit be approximated as:[2]

$$\frac{D_R}{D_i} \approx 1 + \frac{\tau_i^{(\mathrm{ST})}\left(\bar{N}_{\mathrm{ST}}^{(i)}\right)^2}{\tau_0 N^2} \tag{S8}$$

, with $\tau_i^{(\mathrm{ST})}$ the residence time in a complex, to be defined below, and $\bar{N}_{\mathrm{ST}}^{(i)}$ the expected number of non-covalent crosslinks per chain of species $i$. $\tau_0$ is the (non-sticky) monomeric relaxation time and $N$ is the total number of monomers per chain. In our case we have: $\bar{N}_{\mathrm{ST}}^{(i)} = p_i \xi_i$, with $\xi_i$ the valency of species $i$ and $p_i$ the probability that a sticker of a chain of species $i$ is in a bound state. Hence, within our model $\bar{N}_{\mathrm{ST}}^{(i)}$ is not necessarily an integer. Equation S8 shows that the diffusivity ratio depends on the polymer concentration on account of the fact that $p_i$ is concentration dependent. Our calculations assume ideal chains, so that the non-sticky Rouse diffusivity can be expressed as:

$$D_R = \frac{R^2}{\tau_\mathrm{R}} = \frac{b^2 N}{\tau_0 N^2} = \frac{b^2}{\tau_0 N} \tag{S9}$$

, with $\tau_\mathrm{R}$ the Rouse time. Substitution in Equation S9 and rearranging gives:

$$D_i = \frac{b^2}{\tau_0 N}\left[1 + \frac{\tau_i^{(\mathrm{ST})}(p_i \xi_i)^2}{\tau_0 N^2}\right]^{-1} \tag{S10}$$

We now express the diffusivities in terms of the relevant monomeric friction coefficients, taking into account the weighting of sticky and non-sticky monomers:

$$D_R = k_B T / \zeta_0 N \tag{S11}$$

$$D_i = k_B T \Big/ \left[\zeta_0\left(N - \bar{N}_{\mathrm{ST}}^{(i)}\right) + \zeta_{\mathrm{ST}}^{(i)} \bar{N}_{\mathrm{ST}}^{(i)}\right] \tag{S12}$$

, with $\zeta_0$ and $\zeta_{\mathrm{ST}}^{(i)}$ the friction coefficients of non-sticky and sticky monomers, as explained in main text Section 2.4. Combining Equations S9-S12 yields:

$$\frac{D_R}{D_i} = 1 + \frac{\left(\zeta_{\mathrm{ST}}^{(i)} - \zeta_0\right)\bar{N}_{\mathrm{ST}}^{(i)}}{\zeta_0 N} \tag{S13}$$

and

$$\zeta_{\mathrm{ST}}^{(i)} = \zeta_0 \left(1 + \frac{\tau_i^{(\mathrm{ST})} p_i \xi_i}{\tau_0 N}\right) \tag{S14}$$

The residence time $\tau_i^{(\mathrm{ST})}$ is further specified in Section S4.

**3.2. Rouse matrix and viscoelastic moduli**

The relaxation times of the $N-1$ normal modes $\wp$ of a polymer species $i$ containing $N$ segments are given as:[3] $\tau_{i,\wp} = 1/2\lambda_{i,\wp}$, with $\lambda_{i,\wp}$ the eigenvalues of the tridiagonal connectivity matrix $\mathbf{Q}^{(i)}$, of which the non-zero elements are given by:

$$Q_{m,m-1}^{(i)} = -k_{m-1}^{(i)}/\zeta_m^{(i)} \tag{S15}$$

$$Q_{m,m}^{(i)} = k_m^{(i)}/\zeta_m^{(i)} + k_m/\zeta_{m+1}^{(i)} \tag{S16}$$

$$Q_{m,m+1}^{(i)} = -k_{m+1}^{(i)}/\zeta_{m+1}^{(i)} \tag{S17}$$

, with $k_m^{(i)} = k_{m-1}^{(i)} = k_{m+1}^{(i)} = k_0 = 3k_bT/b^2$ the, for simplicity presumed constant, spring constant of a bond between two adjacent monomers and $\zeta_m^{(i)}$ the monomeric friction coefficients. The index $m$ counts the monomers along a chain. Once the connectivity matrices for all species in the droplet or solution have been collected, the sets of eigenvalues are calculated using standard mathematical procedures to give the relaxation times $\{\tau_{i,\wp}\}$ of the normal modes. According to the Boltzmann superposition principle, in the linear viscoelastic regime the storage and loss moduli of the IDP solution can now be obtained by summing the contributions of all chain types/valencies according to a generalized Maxwell model:[4], [5]

$$G'(\omega) = \frac{k_BT}{b^3N}\sum_{i=1}^{n}\left[\phi_i \sum_{\wp=1}^{N-1}\left(\frac{\omega^2\tau_{i,\wp}^2}{1+\omega^2\tau_{i,\wp}^2}\right)\right] \tag{S18}$$

$$\tilde{G}''(\omega) = G''(\omega) - \eta_S\omega = \frac{k_BT}{b^3N}\sum_{i=1}^{n}\left[\phi_i \sum_{\wp=1}^{N-1}\left(\frac{\omega\tau_{i,\wp}}{1+\omega^2\tau_{i,\wp}^2}\right)\right] \tag{S19}$$

, where the latter represents the loss modulus relative to the viscous dissipation by the solvent $\eta_S = \zeta_0/b$.[6] The complex modulus and viscosity are defined as $G^*(\omega) = \sqrt{G'(\omega)^2 + G''(\omega)^2}$ and $\eta^*(\omega) = G^*(\omega)/\omega$. In the Newtonian limit $\omega \to 0$ the terminal viscosity is obtained as: $\eta = \lim_{\omega\to 0} G'(\omega)/\omega$. We note that the sticky Rouse model itself is sequence specific and principally capable of predicting the viscoelastic properties of the solution as a function of the position of the stickers along the IDP backbone.

**Section S4. Effective strand length and sticker life time renormalization**

The friction coefficients $\zeta_{\mathrm{ST}}^{(i)}$ are the same for all stickers of species $i$ as long as there is frequent exchange between binding partners, *i.e.* typically for open networks with a low crosslink density where stickers have considerable probing volume as determined by large strand lengths. As mentioned in the main text, Semenov and Rubinstein argued that in case of a dense network, the generally small strand length suppresses a sticker's probing volume, which increases its effective residence time in a complex due to repeated un- and rebinding before one finds a new binding partner.[7] Since sticky monomers are not necessarily spaced regularly (see main text), this might mean that, due to this renormalization, their friction coefficients not only vary with valency but even *within one and the same chain*.

The minimum strand length $\mathcal{L}_{ij}^{(0)}$ of a sticker $j$ of each a species $i$ with valency $\xi_i = \xi_1 + i - 1$ is taken to be the number of strands that it separates from its nearest neighboring sticker. A strand is a chain section with a length $L_{\mathrm{prec}}$ defined as the (average) number of monomers between evenly spaced and non-terminal precursor units. So, each chain comprises $\xi_n + 1$ such strands with $\xi_n$ the number of precursor sites. For a sticker binding probability $p_i$ < 1, a chain of species $i$ may be in a fully bound, partially bound or even unbound state in equilibrium. In this case, the effective strand length $\mathcal{L}_{ij}$ increases from $\mathcal{L}_{ij}^{(0)}$. We use an approximate method to estimate this increase. The probability of encountering a chain of species $i$ with $0 \leq \xi^{(b)} \leq \xi_i$ of its stickers in a bound state is given by:

$$P\left(\xi^{(b)}\right) = p_i^{\xi^{(b)}}\left(1 - p_{\mathrm{tot}}^{(i)}\right)^{\xi_i - \xi^{(b)}} \binom{\xi^{(b)}}{\xi_i} \tag{S20}$$

, with

$$\sum_{\xi^{(b)}=0}^{\xi_i} P\left(\xi^{(b)}\right) = 1 \tag{S21}$$

Only for $\xi^{(b)} = \xi_i$ the effective strand length is equal to the minimum integer value $\mathcal{L}_{ij} = \mathcal{L}_{ij}^{(0)}$. In the other extreme, for $\xi^{(b)} = 0$, the effective strand length equals $\mathcal{L}_{ij} = \xi_n + 1$, *i.e.* the maximum number of strands per chain. We calculate the effective strand length using the following weighted interpolation:

$$\mathcal{L}_{ij} = \sum_{\xi^{(b)}=0}^{\xi_i} \left[\xi_i^{-1} P\left(\xi^{(b)}\right)\left(\xi^{(b)}\mathcal{L}_{ij}^{(0)} + \left(\xi_i - \xi^{(b)}\right)(\xi_n + 1)\right)\right] \tag{S22}$$

In this model, the sequence specificity is retained only in the minimum strand length $\mathcal{L}_{ij}^{(0)}$ and not in the renormalization that accounts for partially bound states. This is nevertheless acceptable, since for high $p_i$ the deviation from $\mathcal{L}_{ij}^{(0)}$ is small and the loss in specificity hence negligible, whereas for low $p_i$ the interaction between chains is anyway weak. In case of a dense network, the sticker life time renormalizes on account of repetitive binding and unbinding with the same partner. In this case, the residence time of the $j^{\text{th}}$ sticker on a chain of species $i$ in a binary complex becomes: $\tau_{ij}^{(\mathrm{ST})} = \tau_{\mathrm{ST}}^{(0)} \mathcal{L}_{ij}^{-3/2} \phi_A^{-1}$, with $\phi_A$ the volume fraction of free stickers in solution or in a droplet, *i.e.* the mean-field probability of encountering a free sticker. We hence implement the following renormalization for the effective friction coefficient: $\zeta_{ij}^{(\mathrm{ST})} = \zeta_{\mathrm{ST}}^{(i)} \max\left(\tau_{\mathrm{ST}}^{(0)}, \tau_{ij}^{(\mathrm{ST})}\right)/\tau_{\mathrm{ST}}^{(0)}$.

**Section S5. Terminal viscosity**

Figure S2 shows the complex viscosities plotted as a function of frequency. These plots correspond to the calculations behind main text Figure 14.

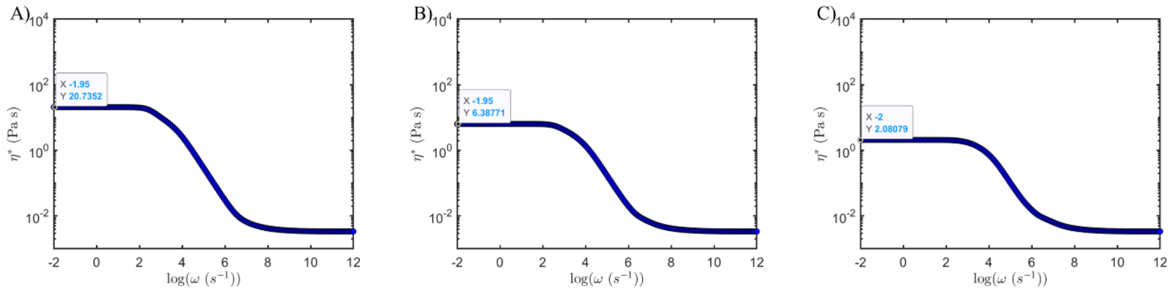


**Figure S2.** Complex viscosity plotted as a function of as a function of frequency for the heptavalent IDP depicted in main text Figure 12, calculated for a dimensionless sticker formation penalty of $\Delta\omega_0$ = 1.10×10$^{-2}$, 1.12×10$^{-2}$ and 1.20×10$^{-2}$ (A to C), with $\chi_{iS}$ = 0.84, $K$ = 16384 M$^{-1}$, $\xi_1$ = 2 and $n$ = 6. The tags report the (approximate) terminal viscosities given in Pa·s.

**Section S6. Input parameters**

Tables S2, S3 and S4 list all input parameters of the model, as well as their description, dimension and evaluation, referring to specific figures and calculations in the main document.

**Table S2.** Flory-Huggins parameters

| parameter | description | dimension | value |
|---|---|---|---|
| $T$ | absolute temperature | K | 290 |
| $N = N_i$ | normalized polymer size | - | 250 |
| $N_S$ | normalized monomer/solvent size | - | 1 |
| $\chi_{ij}$ | monomer-monomer interaction parameter | - | 0 |

| | | | |
|---|---|---|---|
| $\chi_{is}(T) = A + \frac{B}{T}$ | monomer-solvent interaction parameter | - | $A$ = 0; $B$ = 243.0 K [a)]<br>$A$ = 5.0397; B = 1288.5 K [b)]<br>$A$ = 0; $B$ = 163.8 K [c)]<br>$A$ = 2.5198; $B$ = 644.24 K [d)]<br>$A$ = 0; $B$ = 196.56 K [e)]<br>$A$ = 0; $B$ = 327.6 K [f)]<br>$A$ = 0; $B$ = 188.4 K [g)] |
| $\Phi$ | total IDP volume fraction | - | 0.373[h)]; 0.565[i)]; 0.749[j)]; |

[a)]Figure 3A, 8, 10B, 11B, 14, 15; [b)]Figure 3B; [c)]Figure 3C, 4C; [d)]Figure 3D; [e)]Figure 4A, 5A, 5; [f)]Figure 5B; [g)]Figure 4B; [h)]Figure 5A, 6; [i)]Figure 8, 10B, 11B, 14, 15; [j)]Figure 5B.

For the LCST behavior in Figure 3B and 3D, we based our parametrization for $A$ and $B$ on the van 't Hoff plots presented by Görlich et al. on the LCST phase behavior of the perfectly repeated FG-domain prf.GLFG$_{52x12}$.[8] The authors of this work plotted the partitioning coefficient $\ln\left(\frac{C_{\text{dilute}}}{C_{\text{dense}}}\right) = \Delta G$ against $1/T$, with $\Delta G$ the change in the molar Gibbs free energy of partitioning between the coexisting phases. The molar entropy and enthalpy changes, $\Delta S$ and $\Delta H$, can be obtained by the abscissa and slope of the van 't Hoff plot, respectively written as: $-\Delta S/R$ and $\Delta H/R$. We now assume: $A \approx x\,\Delta S/R$ and $B \approx x\,\Delta H/R$, with $x$ << 1 to convert to a per monomer basis. This, together with the values for $\Delta S$ and $\Delta H$ reported in ref. [8], neglecting a moderate dependence on the concentration of background salt, arrives at the numbers used in Figure 3B and 3D (Table S2) for $x$ = 0.0833 and 0.0417, respectively, which, together with the value of $K$ = 16384 M-1 places the relevant transitions in a biological temperature range.

**Table S3.** Parameters relating to sticker valency, formation and association

| parameter | description | dimension | Value |
|---|---|---|---|
| $\xi_1$ | minimum active sticker valency | - | 2[a)]; 3, 4, 5[b)] |
| $n$ | maximum number of IDP species | - | 4[c)]; 6[d)]; 7[e)]; 8[f)]; 9, 10, 11, 12[g)] |
| $\Delta\tilde{u}$ | energy of complexation | K | -2300.13[h)]; -2903.16[i)]; -3305.19[j)] |
| $\Delta\tilde{s}$ | entropy of complexation | - | -1.0[k)] |

| | | | |
|---|---|---|---|
| $K = K_{ij}$ | association constant | l / mol | 1024[l]; 3072[m]; 16384[n]; 524288[o] |
| $\Delta\omega_0$ | sticker formation energy | - | $3.2\times10^{-3}$ [p]; $1.10\times10^{-2}$ – $1.20\times10^{-2}$ [q]; $1.72\times10^{-2}$ – $1.88\times10^{-2}$ [r]; $2.50\times10^{-2}$ [s] |

[a]Figure 3, 4, 5, 6, 8, 10B, 11B, 14, 15; [b]Figure 7; [c]Figure 4, 5, 6, 7; [d]Figure 14, 15; [e]Figure 8, 11B; [f]Figure 10B; [g]Figure 10B, 11B; [h]Figure 4A; [i]Figure 4B; [j]Figure 3, 4C; [k]Figure 3, 4; [l]Figure 5, 6, 7; [m]Figure 10B, 11B; [n]Figure 14, 15; [o]Figure 8; [p]Figure 5, 6, 7; [q]Figure 14, 15; [r]Figure 8; [s]Figure 10B, 11B.

**Table S4.** Parameters relating to dynamics

| parameter | description | dimension | value |
|---|---|---|---|
| $\Gamma_0$ | Effective Arrhenius prefactor for reaction mobility | $s^{-1}$ | Divided out as explained in Section 2.2 |
| $\varepsilon_a^{(f)}$ | Effective activation energy for reaction mobility | in $k_B T$ | Divided out as explained in Section 2.2 |
| $D_0$ | monomer and solvent self-diffusivity | $m^2/s$ | $1.0\times10^{-9}$ |
| $\varepsilon_a = \varepsilon_a^{(ij)}$ | activation energy for binding | in $k_B T$ | 3.45 |

### Section S7. In-vitro phase separation and aging essays of perfect repeat nucleoporin constructs

Using fluorescence recovery after photobleaching (FRAP), we measured aging in phase-separated droplets formed by the following partially Alexa Fluor 488-labeled GLFG-repeat domain constructs, derived from disordered Nup98 FG-repeat domain:[9] GLFG-WT (wildtype), GLFG-$V_8$ and GLFG-$V_{18}$ (construct sequences in Table S5). Phase-separated droplets for each of these variants (examples in Figure S3) formed instantly via rapid buffer exchange procedure described previously.[10] Briefly, purified and labelled GLFG-repeat domain protein was mixed with unlabelled sample in 2M GdmCl and 50mM Tris-HCl, pH8. From this mixture 1µL was then quickly mixed with TB buffer, in a chambered coverslip. For these *in vitro* droplet/FRAP assays, aging time was counted from the moment of the buffer exchange and instantaneous phase-separation. We recorded the FRAP 1h, 24h, 48h and 72h after buffer exchange on multiple droplets within one sample and averaged the results, taking three or two samples per time point per construct. We confirmed reproducibility by measuring at least two different replicates, per GLFG-repeat domain sample and per time interval.

**Table S5.** GLFG-based Nup constructs used for model validation.

| | |
|---|---|
| GLFG-WT | GGLFGGNTQPATGGLFGGNTQPATGGLFGGNTQPATGGLFGGNTQP<br>ATGGLFGGNTQPATGGLFGGNTQPATGGLFGGNTQPATGGLFGGNT<br>QPATGGLFGGNTQPATGGLFGGNTQPATGGLFGGNTQPATGGLFGG<br>NTQPATGGLFGGNTQPATGGLFGGNTQPATGGLFGGNTQPATGGLF<br>GGNTQPATGGLFGGNTQPATGGLFGGNTQPATGGLFGGNTQPATGG<br>LFGGNTQPATGGLFGGNTQPATGGLFGGNTQPATGGLFGGNTQPAT<br>GGLFGGNTQPATGGLFGGNTQPATGGLFGGNTQPATGGLFGGNTQP<br>ATGGLFGGNTQPATGGLFGGNTQPATGGLFGGNTQPATGGLFGGNT<br>QPATGGLFGGNTQPATGGLFGGNTQPATGGLFGGNTQPATGGLFGG<br>NTQPATGGLFGGNTQPATGGLFGGNTQPATGGLFGGNTQPATGGLF<br>GGNTQPATGGLFGGNTQPATGGLFGGNTQPATGGLFGGNTQPATGG<br>LFGGNTQPATGGLFGGNTQPATGGLFGGNTQPATGGLFGGNTQPAT<br>GGLFGGNTQPATGGLFGGNTQPATGGLFGGNTQPATGGLFGGNTQP<br>ATGGLFGGNTQPATGGLFGGNTQPAT* |
| GLFG-V$_8$ | GGLFGGNTQPATGGLFGGNTQPATGGLFGGNTQPATGGLFGGNTQP<br>ATGGLFGGNTQVATGGLFGGNTQPATGGLFGGNTQPATGGLFGGNT<br>QPATGGLFGGNTQPATGGLFGGNTQPATGGLFGGNTQVATGGLFGG<br>NTQPATGGLFGGNTQPATGGLFGGNTQPATGGLFGGNTQPATGGLF<br>GGNTQPATGGLFGGNTQVATGGLFGGNTQPATGGLFGGNTQPATG<br>GLFGGNTQPATGGLFGGNTQPATGGLFGGNTQPATGGLFGGNTQVA<br>TGGLFGGNTQPATGGLFGGNTQPATGGLFGGNTQPATGGLFGGNTQ<br>PATGGLFGGNTQPATGGLFGGNTQVATGGLFGGNTQPATGGLFGGN<br>TQPATGGLFGGNTQPATGGLFGGNTQPATGGLFGGNTQPATGGLFG<br>GNTQVATGGLFGGNTQPATGGLFGGNTQPATGGLFGGNTQPATGGL<br>FGGNTQPATGGLFGGNTQPATGGLFGGNTQVATGGLFGGNTQPATG<br>GLFGGNTQPATGGLFGGNTQPATGGLFGGNTQPATGGLFGGNTQPA<br>TGGLFGGNTQVATGGLFGGNTQPATGGLFGGNTQPATGGLFGGNTQ<br>PATGGLFGGNTQPATGGLFGGNTQPAT* |
| GLFG-V$_{18}$ | GGLFGGNTQVATGGLFGGNTQPATGGLFGGNTQPATGGLFGGNTQ<br>VATGGLFGGNTQPATGGLFGGNTQPATGGLFGGNTQVATGGLFGG<br>NTQPATGGLFGGNTQPATGGLFGGNTQVATGGLFGGNTQPATGGLF<br>GGNTQPATGGLFGGNTQVATGGLFGGNTQPATGGLFGGNTQPATG<br>GLFGGNTQVATGGLFGGNTQPATGGLFGGNTQPATGGLFGGNTQV<br>ATGGLFGGNTQPATGGLFGGNTQPATGGLFGGNTQVATGGLFGGNT<br>QPATGGLFGGNTQPATGGLFGGNTQVATGGLFGGNTQPATGGLFGG<br>NTQPATGGLFGGNTQVATGGLFGGNTQPATGGLFGGNTQPATGGLF<br>GGNTQVATGGLFGGNTQPATGGLFGGNTQPATGGLFGGNTQVATG<br>GLFGGNTQPATGGLFGGNTQPATGGLFGGNTQVATGGLFGGNTQPA<br>TGGLFGGNTQPATGGLFGGNTQVATGGLFGGNTQPATGGLFGGNTQ<br>PATGGLFGGNTQVATGGLFGGNTQPATGGLFGGNTQPATGGLFGGN |

| | |
|---|---|
| | TQVATGGLFGGNTQPATGGLFGGNTQPATGGLFGGNTQVATGGLFG GNTQPATGGLFGGNTQPATGGLFGGNTQVAT* |

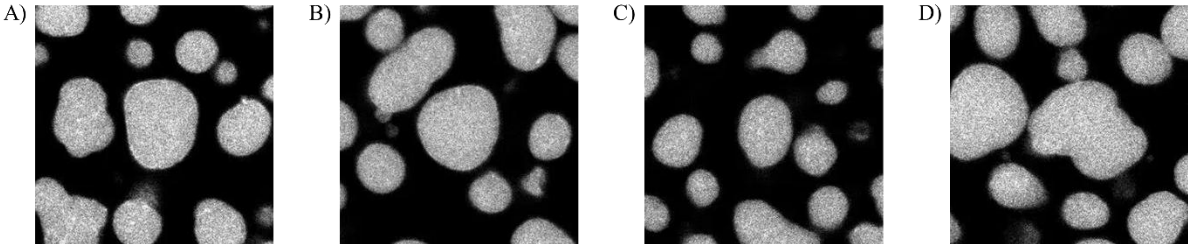


**Figure S3.** Phase separated Nup-rich droplets of GLFG-$V_{18}$ after 1h, 24h, 48h and 72h of aging time.

All normalized FRAP traces were fitted against a biexponential function (see main text Figure 9), from which we obtained the values for $t_{1/2}$. The biexponential function used for the fitting expresses the normalized relative fluorescence intensity $0 \leq \bar{I}(t) \leq 1$ as a function of time $t$ as:

$$\bar{I}(t) = \phi_{\mathrm{MF}} \left\{ w_1 \exp\left[1 - \exp\left(\frac{t - t_{\mathrm{shift}}}{\tau_1}\right)\right] + w_2 \exp\left[1 - \exp\left(\frac{t - t_{\mathrm{shift}}}{\tau_2}\right)\right]\right\} \quad \text{(S21)}$$

, with $\phi_{\mathrm{MF}}$ the mobile fractions, $w_1$ and $w_2$ weights, $\tau_1$ and $\tau_2$ characteristic times and $t_{\mathrm{shift}} \approx$ 10s the time duration of the bleach. During fitting, the mobile fraction $\phi_{\mathrm{MF}}$ were fixed at $\phi_{\mathrm{MF}}$ = 1 for all traces recorded for GLFG-WT and GLFG-$V_8$ at all times, as well as for those recorded for GLFG-$V_{18}$ after 1h and 24h. This choice is supported by the saturation or clear approach to saturation of the recovery curves of GLFG-WT and GLFG-$V_8$, as well as the short-time (1h) aged GLFG-$V_{18}$ at/towards the maximum fractional fluorescence intensity of 1. For this reason, we also assumed a high mobile fractions, close to 1, for the longer aged droplets of GLFG-$V_{18}$. The validity of this choice is strongly supported by the satisfactory fits: for the traces for GLFG-V18 recorded after 48h and 72h, we assumed $\phi_{\mathrm{MF}}$ = 0.85. The values for $w_1$ , $w_2$, $\tau_1$ and $\tau_2$ follow from the fitting and are not further discussed, as we refrain from drawing detailed conclusions from them. What matters are the values for $t_{1/2}$, obtained from individual experimental repeats by solving

$$\frac{1}{2} - \left\{ w_1 \exp\left[1 - \exp\left(\frac{t_{1/2} - t_{\mathrm{shift}}}{\tau_1}\right)\right] + w_2 \exp\left[1 - \exp\left(\frac{t_{1/2} - t_{\mathrm{shift}}}{\tau_2}\right)\right]\right\} = 0 \quad \text{(S22)}$$

The results, averaged over the individual measurements, are listed in Table S6.

**Table S6.** Half times of fluorescence recovery ($t_{1/2}$) of GLFG-repeat domain variants as a function of aging time and valency, given in seconds and averaged over several repeats (see man text, Section 2.4).

| | GLFG-WT | GLFG-$V_8$ | GLFG-$V_{18}$ |
|---|---|---|---|
| 1h | 42 | 75 | 66 |
| 24h | 37 | 85 | 273 |
| 48h | 43 | 77 | 1426 |
| 72h | 42 | 83 | 1357 |

**Section S8. Dynamic arrest**

The way the model is constructed model provides for an approximate description of dynamic arrest or 'solidification'. We will first present the approach, show that, depending on binding strength, the aging kinetics indeed becomes frustrated and then discuss some caveats which are inherent to the model's underlying assumptions and for which reason we treat this exercise as very preliminary. We propose that the decrease in the average polymer diffusivity, defined as $\bar{D}(t) = \frac{\sum_i \phi_i D_i}{\sum_i \phi_i}$, represents a general measure for the frustration of the dynamic processes involved with the aging, *i.e.* sticker formation and dissolution through local folding and unfolding, as well as the binding and unbinding in complexes. We assume that the decrease in diffusivity gives rise to a time-dependent effective contribution to the activation barriers of aforementioned processes:

$$\varepsilon_a^{\mathrm{eff}}(t) = -k_B T \ln\left(\frac{\bar{D}(t)}{\bar{D}_{\mathrm{REF}}}\right) \qquad \text{(S23)}$$

, with $\bar{D}_{\mathrm{REF}} \geq \bar{D}(t)$ a purely phenomenological reference value that determines if and how fast dynamic frustration sets in. In other words, it determines how fast during aging and for a given association strength the system becomes fully frustrated, *i.e.* 'solidifies' out of equilibrium, or eventually still relaxes to equilibrium. We substitute $\varepsilon_a^{\mathrm{eff}}(t)$ in the relations for the reaction mobility for reversible sticker formation, as well as for the bare sticker life time, which now become time-dependent:

$$\Gamma(t) = \Gamma_0 \exp\left[-\beta\left(\varepsilon_a^{(f)} + \varepsilon_a^{\mathrm{eff}}(t)\right)\right] \qquad \text{(S24)}$$

$$\tau_{\mathrm{ST}}^{(0)}(t) = \tau_0 \exp\left[\beta\left(\varepsilon_a + \varepsilon + \varepsilon_a^{\mathrm{eff}}(t)\right)\right] \qquad \text{(S25)}$$

In an iterative calculation, the latter gives an updated average diffusivity, which, through Equation S23 is used to arrive at a new value for the bare complex life time in the next time step.

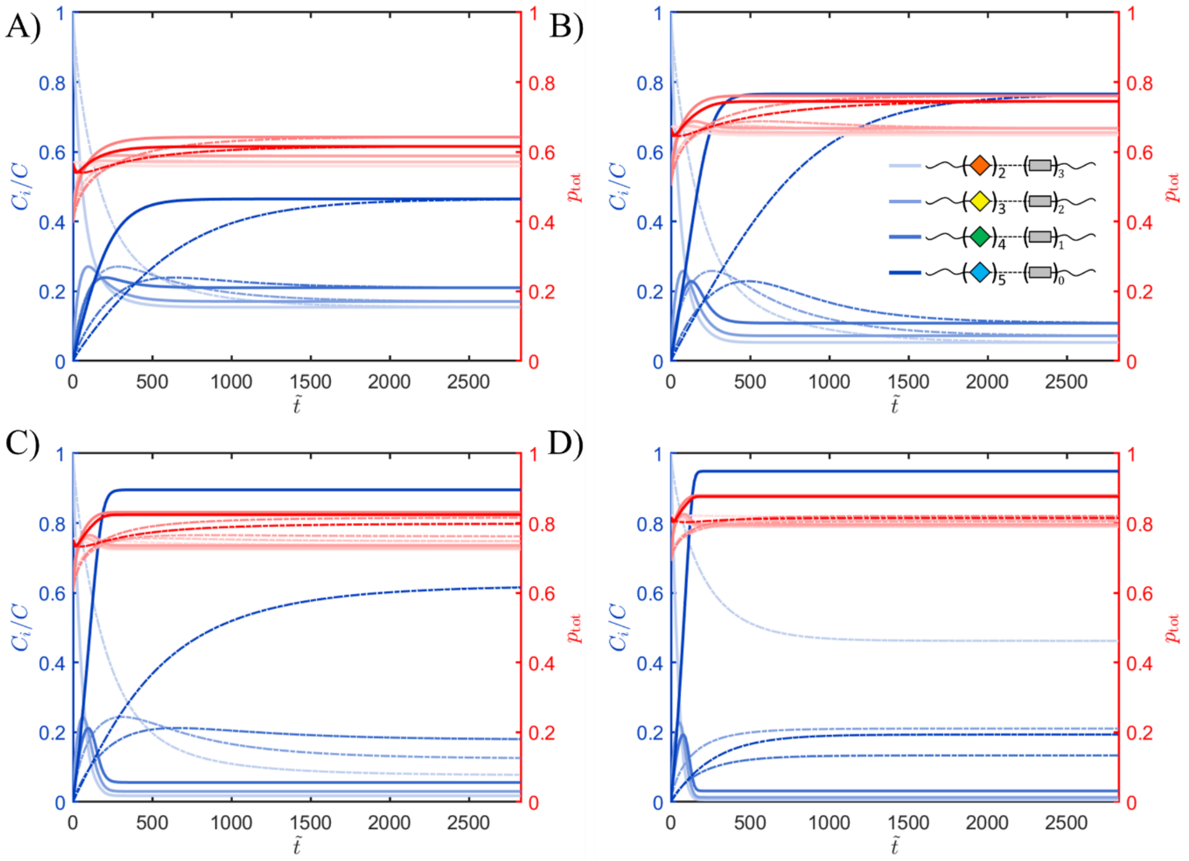


**Figure S4.** Valency distribution as a function of time during kinetic arrest. Panels A) – D) represent traces calculated for a sticker binding constant of $K$ = 512, 10124, 2048 and 4096 $M^{-1}$. All other input parameters are the same as for main text Figure 5A (see Section S6). The plots show the concentration of IDP species $C_i$ with different sticker valencies (see legend in Panel B), normalized by $C_{con}$ (blue curves) and individual (red, shades) and total (red, dark) fractions of bound stickers, all plotted as a function of dimensionless time $\tilde{t} = t\Gamma(\tilde{t} = 0)$. The dash-dotted and solid curves are obtained with and without considering kinetic frustration due to an increasing activation barrier $\varepsilon_a^{\mathrm{eff}}(t)$ (see Equations S28 – S30).

As an example, in Figure S4 we calculate the kinetic traces based on the input parameters underlying main text Figure 5A for four different values for the sticker binding constant and assuming arbitrarily taking $\overline{D}_{\mathrm{REF}} = D_R/\sum_i \phi_i$, *i.e.* the Rouse diffusivity in a semidilute solution under θ-conditions. The traces show that for the present input, a value of $K$ = 512 or 1024 $M^{-1}$ (Figure S4A and S4B) still allows for the system to reach the equilibrium valency distribution, albeit slower than expected based on constant activation barriers. For stronger association, *i.e.* here for $K$ = 2028 and 4096 $M^{-1}$ (Figure S4C and S4D), the traces effectively saturate at fractions away from their equilibrium values, indicating the actual solidification. At this point we would no longer expect the material to behave as a viscoelastic liquid, but rather as a viscoelastic solid.

We end with discussing a few caveats associated with the above description of dynamic arrest. The general description of the arrest seems reasonable. However, $\overline{D}_{\mathrm{REF}}$ is a purely phenomenological parameter and its magnitude has a large effect on whether and when the solution solidifies. Its magnitude is perhaps best determined by comparison with experimental (rheological) on (synthetic) systems for which sticker association strength is known. Our arbitrary choice for $\overline{D}_{\mathrm{REF}}$ to represent the Rouse diffusivity of the non-sticky chain intuitively seems to overestimate the effect of kinetic frustration: one could argue that even despite the multivalency, for pairwise association with strengths of a few thousand $M^{-1}$ (as in Figure S4C and S4D), dynamic arrest would not be expected. This can however be tuned by assuming a different (in this case lower) value for $\overline{D}_{\mathrm{REF}}$. Furthermore, although in dynamic arrest the valency distribution, as expected, ‘freezes’ in an out-of-equilibrium state, in any model formulated according to the original proposed by Semenov and Rubinstein, the bound sticker fraction is always in equilibrium. The latter may not necessarily be guaranteed to be the case in experimental systems.
